# Transposon insertion sequencing of *Pseudomonas aeruginosa* identifies multiple intersecting pathways essential for extreme colistin resistance

**DOI:** 10.64898/2026.04.13.718186

**Authors:** Madeleine B. Vessely, Ryan P. Kich, Samuel W. M. Gatesy, Hanna K. Bertucci, Aliki Valdes, Clara Luczak, Shradha Rao, Artur Muszyński, Parastoo Azadi, Casey N. Kellogg, Brandon L. Jutras, John Mekalanos, Alan R. Hauser, Egon A. Ozer, Kelly E.R. Bachta

## Abstract

Colistin is used to treat antibiotic resistant gram-negative infections, including those caused by *Pseudomonas aeruginosa* (*Pa*). Using a diverse collection of clinical isolates, we identified BWH047, a colistin-resistant isolate with an extremely high minimum inhibitory concentration (MIC, 1280 µg/mL). To characterize the genes conditionally essential for colistin resistance in BWH047, we employed transposon insertion sequencing and identified 20 gene candidates. In-frame deletion validated 75% of the candidates and identified genes in several novel pathways that contribute to colistin resistance, including *algU* and *wapH*. We also identified several candidate genes from previously reported colistin resistance pathways (e.g. *arn*, *pmrAB*). We further investigated the impact of a colistin resistance-associated inner membrane DedA-family undecaprenyl phosphate flippase, which we named DpcA (<u>D</u>edA of *<u>P</u>seudomonas* necessary for <u>c</u>olistin resistance <u>A</u>). Deletion of *dpcA* in BWH047 restored sensitivity to colistin (MIC = 0.5 µg/mL) and resulted in several unique changes to the structure of lipopolysaccharide (LPS), including production of decreased amounts of the colistin resistance-conferring 4-amino-4-deoxy-L-arabinose (L-Ara4N) modification on lipid A. To date, this work represents the most complete analysis of colistin resistance in *Pa* and identifies novel intersecting pathways that contribute to extreme phenotypic resistance.

**Author summary:** *Pseudomonas aeruginosa* is a bacterium that causes a wide variety of infections. It is especially problematic given its propensity to become resistant to antibiotics. One antibiotic used to treat multidrug-resistant *P. aeruginosa* infections is colistin. In this study, we investigated colistin resistance mechanisms in a patient-derived, extremely phenotypically resistant *P. aeruginosa* isolate, BWH047, using transposon insertion sequencing and mass spectrometry. We identified 13 genes conditionally essential for colistin resistance and investigated the role of one of these genes, *dpcA*, on the composition of the bacterial outer membrane, the target of colistin. Additionally, our study identified novel colistin resistance genes residing in several intersecting pathways that could be targeted to prevent the development of antimicrobial resistance.

## Introduction

*Pseudomonas aeruginosa* (*Pa*) is a gram-negative (GN) bacterium that causes a variety of healthcare-associated infections and is often extensively antibiotic resistant. *Pa* quickly acquires resistance to even the newest antimicrobial combinations, forcing clinicians to rely on antibiotics with higher toxicity. Colistin, one such drug, is a cationic antimicrobial peptide (CAMP) with activity against many GN pathogens, including *Pa* [1]. Although it was once sidelined due to toxic side effects, colistin has been re-introduced into clinical use, particularly in combination with other antimicrobials [2–4]. As such, there has been a rapid increase in colistin-resistant infections following its reintroduction [1].

Colistin (also known as polymyxin E), a cationic amphipathic cyclic lipopeptide antibiotic, is believed to work by disrupting lipopolysaccharide (LPS) in the GN outer membrane [5]. Interaction of colistin’s positively charged residues with the negatively charged phosphates on lipid A of LPS is thought to result in displacement of stabilizing divalent cations (e.g. Mg^2+^, Ca^2+^), leading to increased cell permeability and bacterial cell death [5, 6]. To date, the major recognized mechanism of acquired colistin resistance involves modification of lipid A with positively charged moieties that decrease the net negative charge of LPS and consequently reduce colistin’s affinity for the bacterial outer membrane [7, 8]. Lipid A modifications are mediated by sensor-kinase two component systems (TCS) that sense membrane disruption, cation displacement, or the presence of CAMPs and trigger transcription of genes that result in lipid A modification [9].

Although phosphoethanolamine modification of lipid A drastically increases colistin resistance in other bacterial species (e.g. *Escherichia coli, Acinetobacter baumannii,* and *Klebsiella pneumoniae*), this modification has little impact on colistin resistance in *Pa* [10, 11]. In *Pa*, the only lipid A modification known to confer high-level colistin resistance is the addition of 4-amino-4-deoxy-L-arabinose (L-Ara4N) [9, 12]. L-Ara4N synthesis, and its subsequent addition to lipid A, is mediated by the *arn* operon, which is activated by at least five TCS in *Pa* [13–20].

Using a patient-derived, multidrug-resistant isolate of *Pa* with extremely high phenotypic colistin resistance, we identified several new genes conditionally essential for high-level colistin resistance. In addition, we successfully validated well-described resistance mechanisms, lending credence to our approach, and identified novel pathways that contribute to extreme colistin resistance. We specifically examined the role of a newly named gene, *dpcA* (*dpcA* (*<u>d</u>edA* of *<u>P</u>seudomonas* necessary for <u>c</u>olistin resistance <u>A)</u>, in colistin resistance by using mass spectrometry to characterize LPS and lipid A. We demonstrate that deletion of *dpcA* reduces the amount of aminoarabinose modifications found on lipid A, alters the glycosylation and acylation patterns of LPS and lipid A, and leads to colistin sensitivity. These results suggest that lipid A modification remains the critical step for colistin resistance in *Pa*. Our work facilitates a more complete understanding of high level colistin resistance by i) identifying several new resistance pathways, ii) examining the role of *dpcA* in colistin resistance, and iii) providing molecular insights into how we may prevent resistance to critical last resort antibiotics such as colistin and, by proxy, to naturally occurring cationic antimicrobial peptides.

## Results

### Characterization of the PABWH isolate collection

From December 2015 through June 2016 at Brigham and Women’s Hospital in Boston, MA, *Pa* strains were isolated in the clinical microbiology laboratory (n = 100). For this study, 85 isolates (PABWH collection) were chosen representing the first isolate collected from an individual patient (**Fig 1A**). Whole genome sequencing was performed and revealed considerable strain diversity, including 54 distinct sequence types (STs), of which 4 were novel (ST-6655, 6656, 6658, 6659) (**Fig 1A, S4 Table**). Broth microdilution (BMD) was performed in triplicate to establish robust minimum inhibitory concentrations (MICs) for aztreonam, cefepime, ceftazidime, ciprofloxacin, meropenem, piperacillin-tazobactam, and colistin (**Fig 1B, S1 Fig, S4 Table**). Within the PABWH collection, 25.9% of strains were non-susceptible (intermediate or resistant) to aztreonam, 18.8% to cefepime, 15.3% to ceftazidime, 51.8% to ciprofloxacin, 25.9% to meropenem, and 21.2% to piperacillin-tazobactam (**S1 Fig, S4 Table**). In total, four of the isolates (4.7%; PABWH066, PABWH074, PABWH088, PABWH103) met criteria for difficult-to-treat (DTT) *P. aeruginosa* based on the current definition [21].

**Fig 1.**
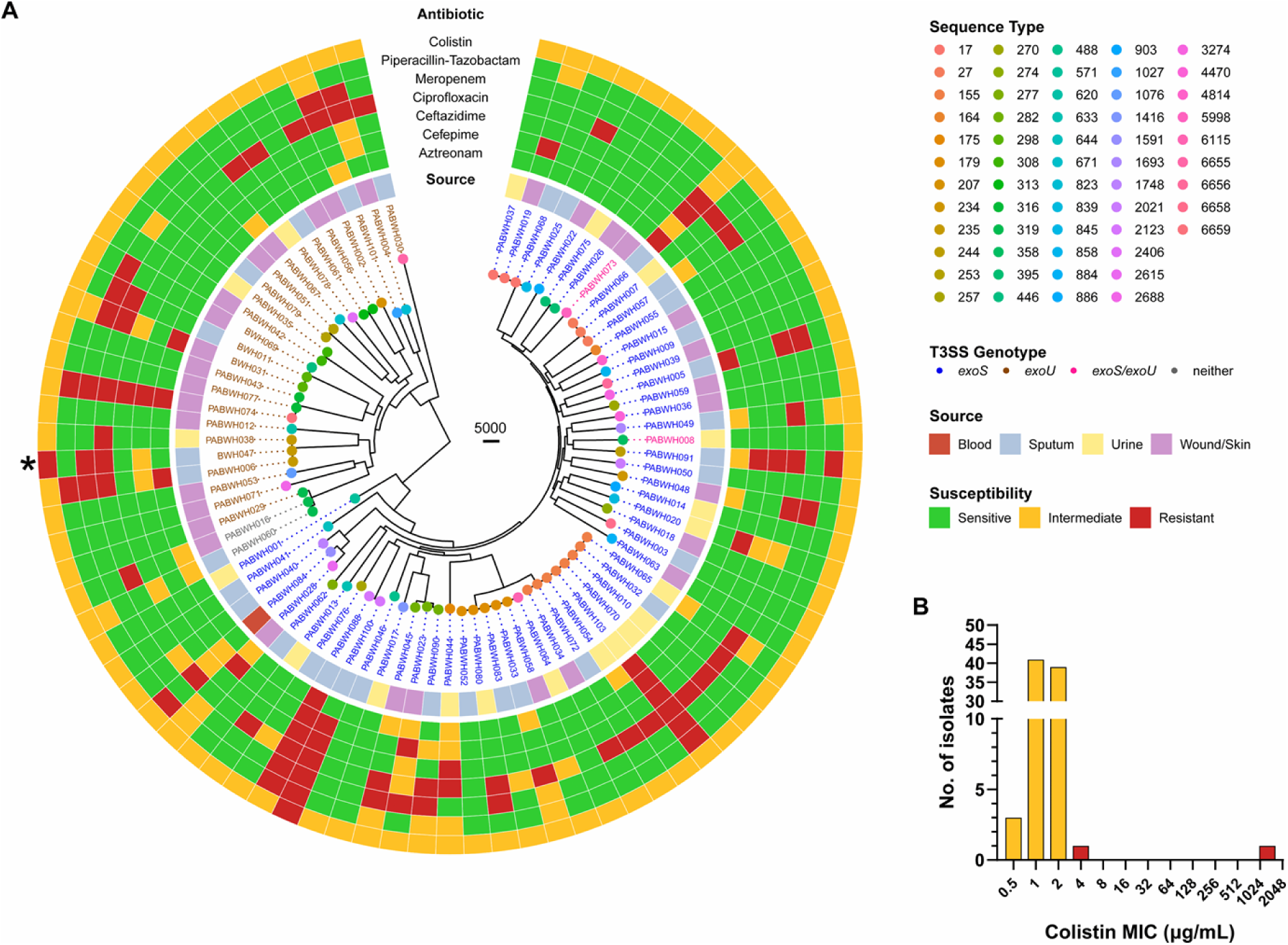
Phylogenetic tree of PABWH isolates and their resistance profiles. (**A**) A maximum likelihood core genome phylogenetic tree was generated with the 85 strains in the PABWH isolate collection. BWH047 is denoted with an asterisk (*). Sequence type is indicated by the colored circles at the tree tips and colored according to legend. Isolate names are colored by Type 3 Secretion genotype (*exoS*, blue; *exoU*, brown; neither, gray; *exoSU,* pink). Source of Isolation is displayed in the inner ring (blood, dark red; sputum, gray; urine, light yellow; wound/skin, purple). Susceptibilities to the indicated antibiotics are displayed in the outer rings (sensitive, green; intermediate, yellow; resistant, red). Susceptibility status was determined according to the CLSI guidelines (M100, Ed. 35) for each antibiotic [25]. (**B**) Colistin MIC (µg/mL), as determined by broth microdilution (BMD), is presented on the x-axis and number of isolates with each MIC is presented on the y-axis. Isolates with intermediate (yellow) and resistant (red) MICs are highlighted. BMD was performed at both standard concentrations and 10x standard concentrations to determine the actual MIC of isolate BWH047.

Isolate BWH047 was a notable outlier within the PABWH collection due to its high colistin MIC (≥128 µg/mL). To determine the exact colistin MIC, we measured MICs at concentrations 10-fold (2.5-1,280 μg/mL) above those typically used for colistin. We found that the actual MIC of BWH047 was 1,280 μg/mL, among the highest ever reported for *Pa* [22–24]. The only other isolate within the PABWH collection that was colistin-resistant had an MIC of 4 μg/mL—the lowest MIC in the ‘resistant’ category according to Clinical and Laboratory Standards Institute (CLSI) guidelines, M100, Ed.35—and 320 times lower than BWH047’s MIC [25] (**Fig 1B**). Of note, in 2019, the CLSI updated the clinical breakpoints for colistin in *Pa* so that there is no longer a susceptible category—a colistin MIC of < 4 μg/mL is categorized as intermediate, while a resistant isolate has an MIC ≥ 4 μg/mL [26]. As such, MICs presented in this paper are interpreted according to the most recent CLSI guidelines [25, 26]. Long-read sequencing analysis of BWH047 revealed that it did not possess any plasmids and the sequence did not contain any of the reported mobile colistin resistance genes *mcr-1-9*. Therefore, we decided to investigate genetic mechanisms driving BWH047’s extreme colistin resistance.

### Identification of genes conditionally essential for extreme colistin resistance in BWH047

To address the question of why BWH047 exhibits such a high colistin MIC, we utilized transposon insertion sequencing (TnSeq). TnSeq is a technique that can be used to identify genes conditionally essential for bacterial survival in the presence of a stressor [27]. We created a BWH047 transposon (tn) library and subsequently exposed it to colistin at ¼ (320 µg/mL) and ½ (640 µg/mL) MICs (**S2C Fig**). Out of 6,424 total genes in the BWH047 genome, we were able to assess the impact of colistin exposure on transposon insertion recovery in 4,157 genes in the 320 µg/mL condition and 4,371 in the 640 µg/mL condition (**S5 & S6 Tables**). For the remainder, we were unable to determine log_2_FC because there were too few reads per gene for analysis. Of note, BWH047 also has a 90,698bp region within the bacterial chromosome at position 2,405,801-2,496,458 bp containing a ∼45kbp duplication. The impact of transposon insertions in this region could not be assessed due to diploidy (**S2B Fig**). Insertion site reads were mapped to the BWH047 genome, and library density and quality were assessed in control (MHB II only) conditions using TIS tools[28]. After filtering, the BWH047:tn library consisted of 16,913 unique transposon insertion sites as defined by ≥ 3 reads in all control pools (n = 5) (**S3 Fig, S5 and S6 Tables**). In total, 27 genes passed our threshold (log_2_FC ≥ −2.0, FDR < 0.05) in 640 μg/mL colistin, and 28 genes met the same criteria in 320 μg/mL colistin (**Fig 2, S7 Table**). Of these, 20 candidate genes were shared in both (**Fig 2, S7 Table**). These genes were deemed conditionally essential genes for BWH047 colistin resistance.

**Fig 2.**
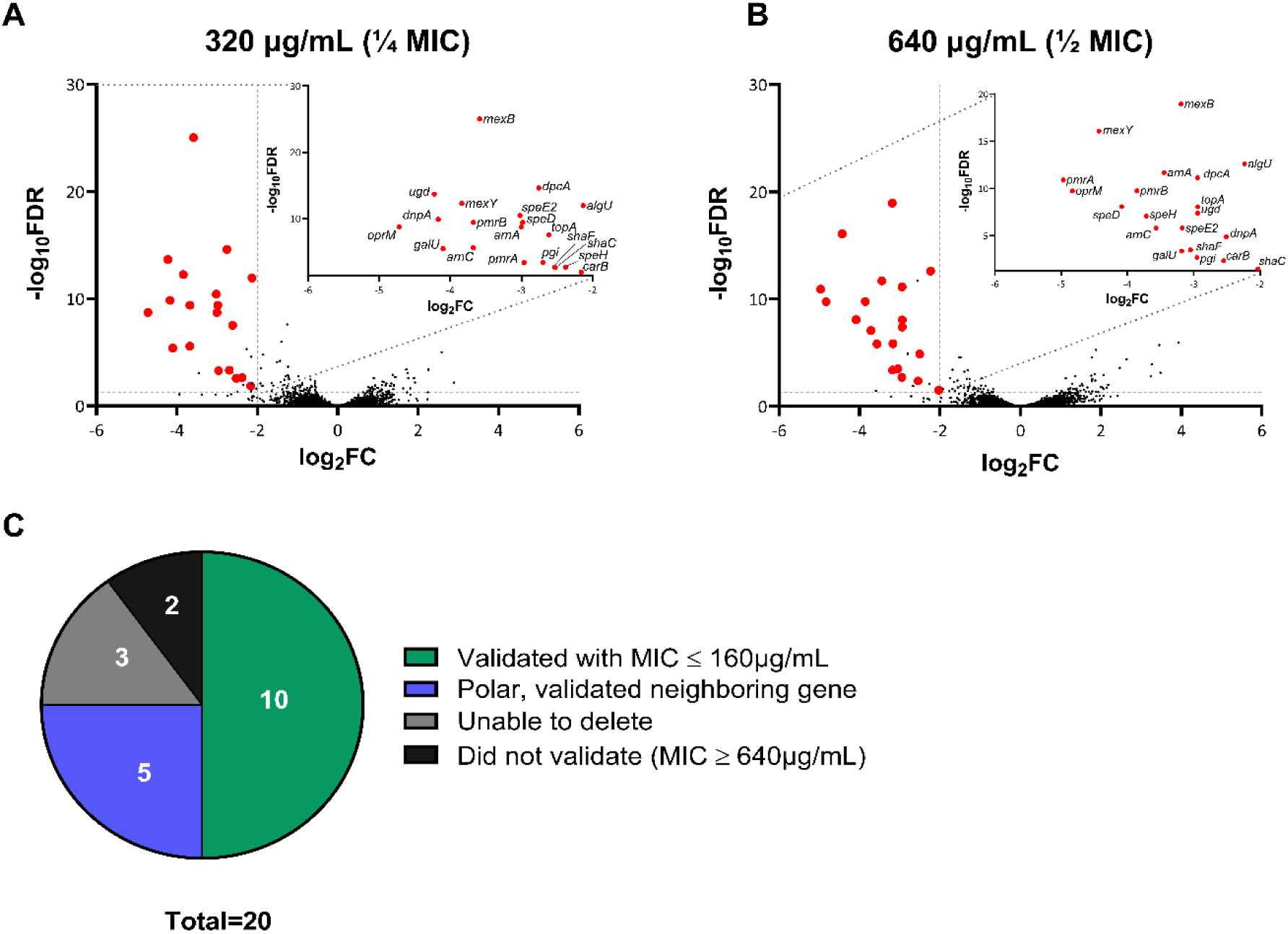
BWH047 genes conditionally essential for colistin resistance. Volcano plots showing TnSeq results following exposure of the BWH047 transposon library to (**A**) 320 and (**B**) 640 μg/mL colistin for 18 hours. The log_2_ fold change (log_2_FC) is presented on the x-axis and the inverse log_10_ of the false discovery rate (FDR) is presented on the y-axis. Red circles indicate conditionally essential genes that passed filtering conditions (log_2_FC ≤ −2.0, -log_10_FDR ≥ 1.3). All 20 overlapping conditionally essential genes are labeled in each upper right hand inset. (**C**) Graphical summary of TnSeq results by target gene validation status.

To validate the TnSeq candidates, we attempted to make in-frame deletions of all of genes on the shared candidate list (**Table 1**, **Fig 2A, B**). We successfully created in-frame deletions in 17 out of 20 genes. For those we were unable to delete (*galU*, *topA*, and *pgi*), we found supporting data to suggest that they may be conditionally essential in certain media types and growth conditions, which may explain why bacteria with transposon insertions in these genes survived during MHBII growth, but we were unable to generate deletions [29, 30]. Additionally, *pgi* and *galU* were previously identified as important for colistin resistance in a resistant *Pa* strain with a colistin MIC of 8 μg/mL, although neither gene target was independently validated [18].

**Table 1.** Colistin minimum inhibitory concentration in deletion mutants and complements of conditionally essential genes identified by TnSeq.

| BWH047 Strain | PAO1 Ortholog | Putative function of gene product | INSeq Hit | Colistin MIC ($\mu\text{g/mL}$) |
| --- | --- | --- | --- | --- |
| wildtype (WT) | - | - |  | 1280 |
| $\Delta\text{mexA}$ | PA0425 | multidrug efflux RND transporter periplasmic adaptor subunit MexA | | 1280 |
| $\Delta\text{mexB}$ | PA0426 | multidrug efflux RND transporter permease subunit MexB | Y | 640 |
| $\Delta\text{oprM}$ | PA0427 | multidrug efflux RND transporter outer membrane channel subunit OprM | Y | 8 |
| $\Delta\text{dpcA}$ | PA4029 | DedA family protein | Y | 0.5 |
| $\Delta\text{dpcA} + \text{dpcA}^a$ | - | Chromosomally complemented $\Delta\text{dpcA}$ strain | | 640 |
| $\Delta\text{algU}$ | PA0762 | RNA polymerase sigma factor AlgU (also known as RpoE) | Y | 0.5 |
| $\Delta\text{shaC}$ | PA1056 | monovalent cation/$\text{H}^+$ antiporter subunit D | Y | 1280 |
| $\Delta\text{shaF}$ | PA1059 | $\text{Na}^+/\text{H}^+$ antiporter subunit G | Y | 640 |
| $\Delta\text{mexY}$ | PA2018 | multidrug efflux RND transporter permease subunit MexY | Y | 8 |
| $\Delta\text{galU}^b$ | PA2023 | UTP-glucose-1-phosphate uridylyltransferase GalU | Y | -- |
| $\Delta\text{topA}^b$ | PA3011 | type I DNA topoisomerase | Y | -- |
| $\Delta arnB$ | PA3552 | UDP-4-amino-4-deoxy-L-arabinose aminotransferase | | 0.5 |
| $\Delta arnC$ | PA3553 | undecaprenyl-phosphate 4-deoxy-4-formamido-L-arabinose transferase | Y | 0.5 |
| $\Delta arnC + arnC^a$ | - | Chromosomally complemented $\Delta arnC$ strain | | 640 |
| $\Delta arnA$ | PA3554 | bifunctional UDP-4-amino-4-deoxy-L-arabinose formyltransferase/UDP-glucuronic acid oxidase ArnA | Y | 0.5 |
| $\Delta arnT$ | PA3556 | lipid IV(A) 4-amino-4-deoxy-L-arabinosyltransferase | | 0.5 |
| $\Delta arnEF^d$ | PA3557, PA3558 | 4-amino-4-deoxy-L-arabinose-phosphoundecapronol flippase subunits ArnE & ArnF | | 0.5 |
| $\Delta ugd$ | PA3559 | UDP-glucose 6-dehydrogenase | Y | 0.5 |
| $\Delta ugd + ugd^a$ | - | Chromosomally complemented $\Delta ugd$ strain | | 640 |
| $\Delta pgi^b$ | PA4732 | glucose-6-phosphate isomerase | Y | -- |
| $\Delta carB$ | PA4756 | carbamoyl-phosphate synthase large subunit | Y | 160 |
| $\Delta speD2$ | PA4773 | adenosylmethionine decarboxylase | Y | 1280 |
| $\Delta speE2$ | PA4774 | spermidine synthase | Y | 1280 |
| $\Delta speH$ | PA4775 | hypothetical protein | Y | 640 |
| $\Delta pmrA$ | PA4776 | two-component system response regulator PmrA | Y | 0.5 |
| $\Delta pmrB$ | PA4777 | two-component system sensor histidine kinase PmrB | Y | 0.5 |
| $\Delta dnpA$ | PA5002 | PIG-L family deacetylase | Y | 640 |
| $\Delta mig14^c$ | PA5003 | antimicrobial resistance protein Mig-14 | | -- |
| $\Delta wapH$ | PA5004 | glycosyltransferase | | 1 |
| $\Delta wapO$ | PA5005 | carbamoyltransferase | | 160 |

<sup>a</sup> $\Delta dpca$ was complemented at the neutral Tn7 site under a constitutive promoter.
$\Delta arnC$ and $\Delta ugd$ were complemented in their native genomic positions under their native promoters.
<sup>b</sup>Though identified as conditionally essential, deletion mutant was unable to be created
<sup>c</sup>We were unable to delete this gene. It was not an original TnSeq candidate but was located in an operon of interest.
<sup>d</sup>MIC for $\Delta arnEF$ were assayed only in the double deletion mutant. Neither gene was identified as conditionally essential by TnSeq analysis.

Once candidate essential gene targets were deleted, all deletion strains were assayed by BMD to determine colistin MIC (**Table 1**). Ten deletion mutants had colistin MICs that were reduced by at least eight-fold compared to BWH047 (parental isolate), including seven which had MICs of 0.5 μg/mL, well below the CLSI threshold for *Pa* colistin resistance and consistent with previously susceptible isolates [25] (**Fig 2C**, **Table 1**). Validated hits included: several members of the *arn* operon (*arnA, arnC, ugd*), the two component system *pmrAB*, the carbamoyl synthesis gene *carB,* the resistance nodulation division (RND) efflux proteins *mexY* and *oprM*, the sigma factor *algU* (otherwise known as *rpoE*), and a DedA-family undecaprenyl phosphate (UndP) carrier protein hereafter designated *dpcA* for <u>d</u>edA of *<u>P</u>seudomonas aeruginosa* required for <u>c</u>olistin resistance [31]. Below, we discuss each of these gene groups in more detail.

Deletion of *arnA*, *arnC*, or *ugd* resulted in a 2,560-fold decrease in colistin MIC from 1280 µg/mL to 0.5 µg/mL (**Table 1**). The *arnBCADTEF-ugd* operon is responsible for the biosynthesis and addition of 4-amino-4-deoxy-L-arabinose (L-Ara4N) to the phosphate moieties on lipid A, and is a well-characterized mechanism of colistin resistance in many bacteria, including *Pa* [18, 31, 32]. Therefore, the identification of multiple genes located within this operon served as an internal positive control that validated our TnSeq experiment. Although we only identified three members of this operon in our experiment, it is well established that all genes in the *arn* operon are necessary for the synthesis and addition of L-4AraN to lipid A [33, 34]. Thus, we investigated the conditional essentiality of additional genes in this operon (*arnB, arnT, arnE, arnF*) and found that all deletions resulted in a uniform colistin MIC of 0.5 μg/mL (**Table 1**). Finally, we restored BWH047’s extreme colistin resistance phenotype through complementation of Δ*arnC* (in-frame deletion) and Δ*ugd* with their native alleles, whereby Δ*arnC* + *arnC* (complementation of *arnC* gene deletion) and Δ*ugd* + *ugd* had colistin MICs of 640 μg/mL, which is within one 2-fold dilution of the parental isolate and within the acceptable range of error of the BMD assay (**Table 1**).

Five two component systems (TCS) that have been shown to induce *arn* expression in *Pa*—PmrAB, PhoPQ, CprRS, ColRS, and ParRS [12, 17–20, 35, 36]. In BWH047, only one of these TCS was identified as conditionally essential for colistin resistance—PmrAB. As with the *arn* operon, deletion of either gene resulted in a 2,560-fold decrease in colistin MIC from 1280 µg/mL to 0.5 µg/mL (**Table 1**).

Additionally, TnSeq identified three candidate targets near *pmrAB* including *speD2, speE2,* and *speH*; however, deletion of any of these three candidates resulted in no change in colistin MIC (**Table 1**). Intriguingly, these three candidate genes are located directly upstream of *pmrAB* in a proposed operon (**S2C Fig**), providing a likely explanation for their identification in the TnSeq experiment despite our inability to validate them. Whether *speD2, speE2, speH, pmrA,* and *pmrB* are all co-transcribed or are expressed as two separate operonic units is unclear [36, 37].

TnSeq also identified conditionally essential targets involved in the modification of LPS core oligosaccharides (core-OS). Deletion of *carB*, a carbamoyl phosphate synthase that works with putative carbamoyl transferase WapO to carbamoylate sugar residues in the core-OS of *Pa[38, 39],* resulted in a colistin MIC of 160 μg/mL, an 8-fold decrease from BWH047. Deletion of *wapO*, though not identified as a candidate in the original TnSeq analysis, resulted a similar MIC (160 µg/mL). One of our non-validating targets, *dnpA* (MIC 1280 µg/mL), was also located within the same operon as *wapO* (*wapO-wapH-mig14-dnpA*). Thus, we elected to investigate the conditional essentiality of the remaining genes in this operon, including *wapH* and *mig14* (**S2C Fig**). To this end, we successfully generated a clean deletion of *wapH*. We were unable to delete the hypothetical *mig14*, which has been previously reported for *Pa* [40]. The colistin MIC of BWH047Δ*wapH* was 1 μg/mL (**Table 1**), which supports the hypothesis that transposon insertions in *dnpA* may have influenced expression of the remaining genes within the same operon. Polar transposon effects may have led to *dnpA*’s identification as conditionally essential when the true conditionally essential genes (*wapH* and *wapO*) were located elsewhere in the same operon.

Both *mexY* and *oprM* were identified as conditionally essential for colistin resistance, and deletion of either gene resulted in a colistin MIC of 8µg/mL. These genes code for members of the MexAB-OprM and MexXY-OprM resistance-nodulation-division (RND) transporters, which are well-characterized tripartite efflux systems in *Pa*. Both extrude a wide variety of antimicrobials and promote antimicrobial resistance; however, neither has been shown to export colistin or polymyxin B [41]. MexY spans the inner membrane and OprM is the outer membrane channel protein that is shared by several efflux systems including both MexAB-OprM and MexXY-OprM [42]. Interestingly, we also identified *mexB* as a candidate; however, deletion of *mexB* resulted in a MIC similar to the parental isolate (640 µg/mL). This suggests that the MexXY-OprM efflux pump, but not the MexAB-OprM efflux pump, contributes to colistin resistance in BWH047. Like *speD2, speE2,* and *speH,* we hypothesize that transposon insertions in *mexB* lead to polar effects impacting the expression of *oprM,* which resides downstream of *mexAB* in a transcriptional unit (**S2C Fig**). Therefore, genetic proximity likely led to erroneous identification of *mexB* as a candidate.

One of the more intriguing candidates identified was the sigma factor *algU,* which, when deleted, resulted in a colistin MIC of 0.5µg/mL. *AlgU*, also known as *rpoE* in most other GN bacteria, encodes the extracellular sigma factor (ECF) σ^22^ in *Pa*, which is most often studied in the context of mucoidy in chronic infections [43, 44].

Next, we identified a DedA-family undecaprenyl phosphate flippase, hereafter named *dpcA*, as conditionally essential for colistin resistance in BWH047. Deletion of *dpcA* also resulted in a colistin MIC of 0.5µg/mL, which could be complemented by expression of *dpcA* at a neutral site in the chromosome (**Table 1**). We renamed this gene *dpcA* for *<u>D</u>edA* of *<u>P</u>seudomonas* necessary for <u>c</u>olistin resistance <u>A</u> according to the precedent set in other GN bacterial genera including *Burkholderia* and *Klebsiella* [31, 45]. In these species, orthologs of *dpcA* are required for colistin resistance [31, 45–47]. We sought to better characterize its function in BWH047.

Finally, we were unable to validate a role in colistin resistance for five experimentally identified candidate genes: *shaC, shaF*, *galU*, *topA*, and *pgi,* either because deletion had no impact on colistin MIC (*shaC*, MIC 1280µg/mL, and *shaF*, MIC 640µg/mL; **Table 1**) or we were unable to generate in-frame deletions (*galU*, *topA*, and *pgi*). After accounting for genomic architecture, the overall validation rate of our experiment increased from 10/20 to 15/20, or 75% (**Fig 2C**), which aligns with similar TnSeq experiments [46, 48–50].

### Lipopolysaccharide (LPS) analysis of BWH047 WT and *Δ*dpcA

Given the enrichment of lipopolysaccharide (LPS) modification genes in our experimental results and the fact that the bacterial outer membrane and its structures serve as the binding target for colistin, we hypothesized that the LPS structure in BWH047 could be contributing to its uniquely elevated colistin MIC. Thus, we decided to characterize the LPS of BWH047 and its colistin-sensitive derivative BWH047Δ*dpcA* and Δ*wapH.* LPS from BWH047, Δ*dpcA,* and Δ*wapH* was isolated utilizing the hot phenol-water method [51] and visualized by silver staining after deoxycholate-polyacrylamide gel electrophoresis (DOC-PAGE) gels. The laddering of O-antigen is reduced in the Δ*dpcA* strain compared to the WT BWH047 strain, consistent with the proposed role of DpcA in UndP recycling and UndP’s involvement in transporting O-antigen subunits [52, 53]. Δ*wapH* exhibited an even more extreme truncated version of LPS lacking any visible O-antigen laddering (**S4B Fig**). LPS of BWH047 and Δ*dpcA* was further hydrolyzed with hydrochloric acid in methanol to determine the glycosyl and fatty acid residues (**S4C and D Fig**). Comparative GC-MS analysis showed that both strains contained glycosyl residues that could be attributed specifically to: (i) the O-antigen of *Pa* (N-acetylfucosamine (FucNAc) and N-acetylquinovosamine (QuiNAc)), (ii) the core-OS (3-deoxy-D-manno-oct-2-ulosonic-acid (Kdo)), and (iii) either in the core-OS or the O-antigen, (N-acetylgalactosamine (GalNac), glucose (Glc), N-acetylglucosamine (GlcNAc), mannose (Man), N-aceytylgalactosamine (GalNAc), and rhamnose (Rha)). Heptose could not be identified in the applied chemical derivatization method, likely due to complete phosphorylation of the residue as previously reported [54].

In addition, we observed differences in the relative molar percentage of glycosyl residues in BWH047 and Δ*dpcA* LPS (**Table 2**). In BWH047Δ*dpcA*, we found a significant decrease in the relative content of FucNAc (50% to 1.2%) and relative increases in Rha (2.3% to 9.6%), Glc (40.7 % to 66.2%), Kdo (2.5% to 14.6%), and GalNAc (0.7% to 3.6%). The relative content of GlcNAc also increased in BWH047Δ*dpcA* (2.7% compared to 0.2%). These findings are all consistent with the decrease of O-antigen laddering observed in BWH047Δ*dpcA* (**S4A and B Fig**). Of note, L-Ara4N was the only glycosyl residue attributed to lipid A that decreased in relative mole percentage in BWH047Δ*dpcA* (1.9%) compared to the parental isolate (3.0%).

**Table 2.** Relative molar percentage of the glycosyl composition of lipopolysaccharide from BWH047 WT and Δ*dpcA* as determined by gas chromatography-mass spectrometry.

| Glycosyl Residue |  | Strain |  |
| --- | --- | --- | --- |
| | | BWH047 | BWH047$\Delta dpcA$ |
| Rhamnose | Rha | 2.3 | 9.6 |
| 4-amino-4-deoxy-L-arabinose<sup>a</sup> | L-Ara4N | 3.0 | 1.9 |
| N-acetyl fucosamine<sup>a</sup> | FucNAc | 50 | 1.2 |
| Mannose | Man | 0.4 | 0.1 |
| Glucose | Glc | 40.7 | 66.2 |
| 3-deoxy-D-manno-oct-2-ulosonic acid | Kdo | 2.5 | 14.6 |
| N-acetylquinovosamiosyl<sup>a</sup> | QuiNAc | 0.1 | tr. |
| N-acetylgalactosamine | GalNac | 0.7 | 3.6 |
| L-glycero-D-manno-heptose | Hep | tr. | tr. |
| N-acetylglucosamine | GlcNac | 0.2 | 2.7 |

<sup>a</sup>Due to the unavailability of authentic standards of FucNac, L-Ara4N, and QuiNac, we used the relative response factor of GlcNac to calculate the relative mole percentage. Trace (tr.) was defined as a relative mole percentage < 0.1%.

We also compared the fatty acid compositions of BWH047 and Δ*dpcA* LPS using GC-MS (**Table 3**, **S4C and D Fig**). The LPS of BWH047 was constituted with 3-hydroxydecanoic acid (10:0(3-OH)), 2-hydroxydodecanoic acid (12:0(2-OH)), 3-hydroxydodecanoic acid (12:0(3-OH)), hexadecanoic acid (palmitic acid, 16:0), and, in addition, cis-8-octadecanoic acid (18:1Δ^8^). Δ*dpcA* showed increased amounts of 2-and 3-hydroxydodecanoic acid compared to BWH047 (21% vs. 15% and 52% vs. 43%, respectively), and a reduced amount of palmitic acid (6% vs. 16%). We did not detect cis-8-octadecanoic acid in Δ*dpcA* LPS. Overall, fatty acid composition analysis of BWH047 and Δ*dpcA* LPS revealed both a high degree of heterogeneity in lipid A acylation status and isolate-specific acylation patterns.

**Table 3.** Comparative fatty acid analysis (FAME) of lipopolysaccharide isolated from BWH047 WT and Δ*dpcA*.

| FAME (TMS)<sup>a</sup> |  | Strain |  |
| --- | --- | --- | --- |
| | | BWH047 | BWH047$\Delta dpcA$ |
| 3-hydroxydecanoic acid | 10:0 (3-OH) | 17 | 18 |
| 2-hydroxydodecanoic acid | 12:0 (2-OH) | 15 | 21 |
| 3-hydroxydodecanoic acid | 12:0 (3-OH) | 43 | 52 |
| dodecanoic (lauric) acid | 12:0 | 4 | 3 |
| hexadecanoic (palmitic) acid | 16:0 | 16 | 6 |
| cis-8-octadecanoic acid | 18:1$\Delta^8$ | 5 | n.d. |

<sup>a</sup>Values presented are calculated based on relative response from the GC-MS EI detector.
Abbreviations: not detected (n.d.)

### Lipid A analysis of BWH047 WT and Δ*dpcA* by MALDI-TOF MS

To better characterize the structure of lipid A and to interrogate levels of L-Ara4N modification and the differences in fatty acid distribution, we released lipid A from LPS with a mild sodium acetate treatment and analyzed using MALDI-TOF MS (Matrix-Assisted Laser Desorption/Ionization Time-of-Flight Mass Spectrometry) (**Fig 3A, B**). For the peaks for which we were able to assign chemical compositions (**Table 4**), we calculated relative percentages of total lipid A with the indicated number of L-Ara4N modifications and acyl chains. A higher percentage of lipid A structures were modified by ≥ 1 aminoarabinose moiety in BWH047 compared to Δ*dpcA* (38.9% vs. 13.4%, respectively) (**Table 5**). Additionally, BWH047 had more penta-acylated lipid A compared to Δ*dpcA* (89.3% vs. 76.5%) and Δ*dpcA* had higher levels of tetra-(12.3% vs. 4.3%) and hexa-acylated lipid A (11.2% vs. 6.4%). Penta-acylation was the most predominant pattern in both WT and Δ*dpcA* (**Table 6**).

**Fig 3.**
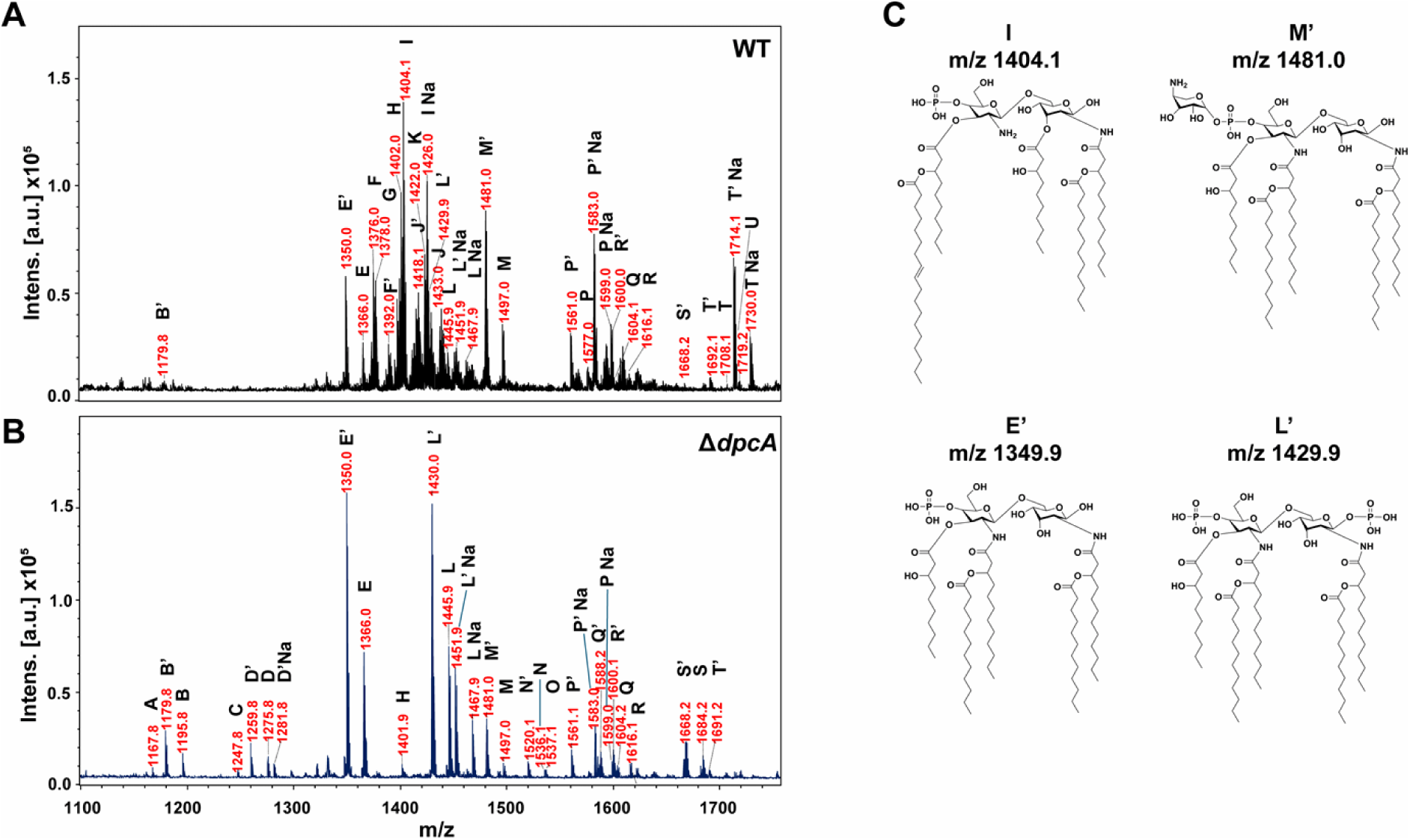
MALDI-TOF analysis of lipid A from BWH047 and Δ*dpcA* strains. Comparative Matrix-Assisted Laser Desorption/Ionization Time-of-Flight (MALDI-TOF) analysis of lipid A from BWH047 (**A**) and BWH047Δ*dpcA* (**B**) released by mild hydrolysis with sodium acetate at pH 4.5. The spectra were acquired in negative ionization mode ([M-H]^-^). The x-axis represents the mass to charge ratio (m/z) and the y-axis represents the intensity of each peak in arbitrary units (a.u.) x 10^5^. (**C**) Structural predictions of selected peaks (identified by letter in panels A and B) differentially present in either BWH047 or Δ*dpcA* and their corresponding predicted m/z values. The detailed assignments are listed in **Table 4** and **S6 Fig**.

**Table 4.** Proposed composition of selected peaks from BWH047 and BWH047Δ*dpcA* lipid A identified by Matrix-Assisted Laser Desorption/Ionization Time-of-Flight (MALDI-TOF) mass spectrometry.

| Label | BWH047 WT | BWH047 $\Delta dpcA$ | [M-H] | Proposed Chemical Composition<sup>a</sup> |
| --- | --- | --- | --- | --- |
| | Observed $m/z$<sup>b</sup> | | Calculated $m/z$ | |
| A |  | 1167.8 | 1167.7 | P, GlcNAc2, 10:0(3-OH), 12:0, [12:0(3-OH)]2 |
| B' | 1179.8 | 1179.8 | 1179.8 | P, GlcNAc2, [12:0]2, [12:0(3-OH)]2 |
| B |  | 1195.8 | 1195.8 | P, GlcNAc2, 12:0(2-OH), [12:0(3-OH)]2, 12:0 |
| C |  | 1247.8 | 1247.7 | P2, GlcNAc2, 10:0(3-OH), 12:0, [12:0(3-OH)]2 |
| D' |  | 1259.8 | 1259.7 | P2, GlcNAc2, [12:0]2, 12:0(3-OH)2 |
| D |  | 1275.8 | 1275.7 | P2, GlcNAc2, 12:0(2-OH), [12:0(3-OH)]2, 12:0 |
| D' Na |  | 1281.8 | 1281.8 | P2, GlcNAc2, [12:0]2, 12:0(3-OH)2 + Na |
| E' | 1350.0 | 1350.0 | 1349.9 | P, GlcNAc2, 10:0(3-OH), [12:0]2, [12:0(3-OH)]2 |
| E | 1366.0 | 1366.0 | 1365.9 | P, GlcNAc2, 10:0(3-OH), 12:0(2-OH), [12:0(3-OH)]2, 12:0 |
| F' | 1376.0 | | 1375.9 | P, GlcNAc2, 18:1$\Delta^8$, [10:0(3-OH)]2, 10:0, 12:0 (3-OH) |
| G | 1378.0 | 1378.0 | 1378.8 | L-Ara4N, P2, GlcNAc2, 10:0(3-OH), 12:0, [12:0(3-OH)]2 |
| F | 1392.0 | | 1391.9 | P, GlcNAc2, 18:1$\Delta^8$, [10:0(3-OH)]2, 10:0(2-OH), 12:0 (3-OH) |
| H | 1402.0 | 1401.9 | 1401.8 | P2, GlcNAc2, [10:0(3-OH)]2, [12:0]2, 12:0(3-OH) |
| I | 1404.1 | 1404.1 | 1404.0 | P, GlcNAc2, 18:1$\Delta^8$, [10:0(3-OH)]2, 12:0, 12:0(3-OH) |
| J' | 1418.1 |  | 1417.8 | P2, GlcNAc2, [10:0(3-OH)]2, (12:0), [12:0(3-OH)]2 |
| K | 1422.0 |  | 1422.0 | P, GlcNAc2, 16:0, 10:0(3-OH), 12:0(2-OH), [12:0(3-OH)]2 |
| I Na | 1426.0 | 1426.1 | 1426.0 | P, GlcNAc2, 18:1$\Delta^8$, [10:0(3-OH)]2, 12:0, 12:0(3-OH) + Na |
| L' | 1429.9 | 1430.0 | 1429.9 | P2, GlcNAc2, 10:0(3-OH), [12:0]2, [12:0(3-OH)]2 |
| J | 1433.0 |  | 1433.8 | P2, GlcNAc2, [10:0(3-OH)]2, [12:0(2-OH)]2, 12:0(3-OH) |
| L | 1445.9 | 1445.9 | 1445.9 | P2, GlcNAc2, 10:0(3-OH), 12:0(2-OH), [12:0(3-OH)]2, 12:0 |
| L' Na | 1451.9 | 1451.9 | 1451.9 | P2, GlcNAc2, 10:0(3-OH), [12:0]2, [12:0(3-OH)]2 + Na |
| L Na | 1467.9 | 1467.9 | 1467.9 | P2, GlcNAc2, 10:0(3-OH), 12:0(2-OH), [12:0(3-OH)]2, 12:0 + Na |
| M' | 1481.0 | 1481.0 | 1481.0 | L-Ara4N, P, GlcNAc2, 10:0(3-OH), [12:0]2, [12:0(3-OH)]2 |
| M | 1497.0 | 1497.0 | 1497.0 | L-Ara4N, P, GlcNAc2, 10:0(3-OH), 12:0(2-OH), [12:0(3-OH)]2, 12:0 |
| N' |  | 1520.1 | 1520.0 | P, GlcNAc2, [10:0(3-OH)]2, [12:0]2, [12:0(3-OH)]2 |
| N |  | 1536.1 | 1536.0 | P, GlcNAc2, [10:0(3-OH)]2, 12:0(2-OH), [12:0(3-OH)]2, 12:0 |
| O |  | 1537.1 | 1537.0 | L-Ara4N, P, GlcNAc2, 16:0, 10:0(3-OH), 12:0, [12:0(3-OH)]2 |
| P' | 1561.0 | 1561.0 | 1560.9 | L-Ara4N, P2, GlcNAc2, 10:0(3-OH), [12:0]2, [12:0(3-OH)]2 |
| P | 1577.0 |  | 1576.9 | L-Ara4N, P2, GlcNAc2, 10:0(3-OH), 12:0(2-OH), [12:0(3-OH)]2, 12:0 |
| P' Na | 1583.0 | 1583.0 | 1582.9 | L-Ara4N, P2, GlcNAc2, 10:0(3-OH), [12:0]2, [12:0(3-OH)]2 + Na |
| Q' |  | 1588.2 | 1588.1 | P, GlcNAc2, 16:0, 10:0(3-OH), (12:0)2, [12:0(3-OH)]2 |
| P Na | 1599.0 | 1599.0 | 1598.9 | L-Ara4N, P2, GlcNAc2, 10:0(3-OH), 12:0(2-OH), [12:0(3-OH)]2, 12:0 + Na |
| R' | 1600.0 | 1600.1 | 1600.0 | P2, GlcNAc2, [10:0(3-OH)]2, (12:0)2, [12:0(3-OH)]2 |
| Q | 1604.1 | 1604.2 | 1604.1 | P, GlcNAc2, 16:0, 10:0(3-OH), 12:0(2-OH), [12:0(3-OH)]2, 12:0 |
| R | 1616.1 | 1616.1 | 1616.0 | P2, GlcNAc2, [10:0(3-OH)]2, 12:0(2-OH), [12:0(3-OH)]2, 12:0 |
| S' | 1668.2 | 1668.2 | 1668.1 | P2, GlcNAc2, 16:0, 10:0(3-OH), [12:0]2, [12:0(3-OH)]2 |
| S |  | 1684.2 | 1684.1 | P2, GlcNAc2, 16:0, 10:0(3-OH), 12:0(2-OH), [12:0(3-OH)]<sub>2</sub>, 12:0 |
| T' | 1692.1 | 1691.2 | 1691.9 | L-Ara4N2, P2, GlcNAc2, 10:0(3-OH), [12:0]<sub>2</sub>, [12:0(3-OH)]<sub>2</sub> |
| T | 1708.1 |  | 1708.0 | L-Ara4N2, P2, GlcNAc2, 10:0(3-OH), 12:0(2-OH), [12:0(3-OH)]<sub>2</sub>, 12:0 |
| T' Na | 1714.1 | 1714.0 | 1714.0 | L-Ara4N2, P2, GlcNAc2, 10:0(3-OH), [12:0]<sub>2</sub>, [12:0(3-OH)]<sub>2</sub> + Na |
| U | 1719.2 | 1719.3 | 1719.2 | L-Ara4N, P1, GlcNAc2, 16:0, 10:0(3-OH), [12:0]<sub>2</sub>, [12:0(3-OH)]<sub>2</sub> |
| T Na | 1730.0 |  | 1730.0 | L-Ara4N2, P2, GlcNAc2, 10:0(3-OH), 12:0(2-OH), [12:0(3-OH)]<sub>2</sub>, 12:0 + Na |

<sup>a</sup>Due to the large number of species present in each strain, we assigned chemical compositions for a selection of peaks. Chemical structures of species with proposed compositions are depicted in Fig 3C and S5 Fig. Raw signal integration values are listed in Supplementary Table 9.
<sup>b</sup>The m/z values obtained in the negative reflector were compared and matched with the calculated monoisotopic ions for the most probable structures.
Abbreviations: phosphate (P), N-acetyl-glucosamine (GlcNAc), 4-amino-4-deoxy-L-arabinose (L-Ara4N), sodium (Na), 3-hydroxidecanoic acid (10:0 (3-OH)), 2-hydroxydodecanoic acid (12:0 (2-OH)), 3-hydroxydodecanoic acid (12:0 (3-OH)), dodecanoic (lauric) acid (12:0), hexadecenoic (palmitic) acid (16:0), cis-8-octadecanoic acid (18:1 $\Delta^8$ )

**Table 5.**
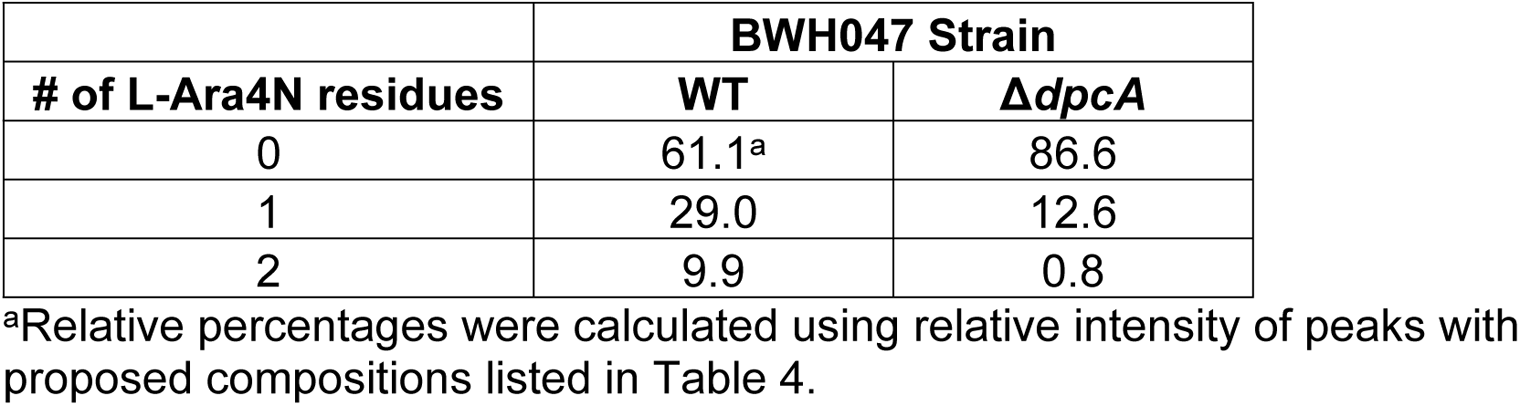
Relative percentage of lipid A structures with aminoarabinose modification(s).

**Table 6.** Relative percentage of lipid A structures with tetra-, penta-, or hexa-acyl chains.

| # of acyl chains | BWH047 Strain |  |
| --- | --- | --- |
| | WT | $\Delta dpcA$ |
| Tetra | 4.3<sup>a</sup> | 12.3 |
| Penta | 89.3 | 76.5 |
| Hexa | 6.4 | 11.2 |

<sup>a</sup>Relative percentages were calculated using relative intensity of peaks with proposed compositions listed in Table 4.

For several prominent peaks in the WT spectrum (m/z 1375.9 (Peak F’), 1391.9 (Peak F), and 1404.4 (Peak I), **Fig 3**), we had difficulty solving structures. We elected to perform collision-induced tandem mass spectrometry (CID MS/MS) analysis of all three selected ions, which pointed to pentaacylated lipid A species substituted with one acyloxy-acyl cis-8-octadecanoic acid (18:1Δ^8^) on C2 of GlcN II (F’, F, I, I Na) (**S5 Fig**). The lipid A species containing 18:1Δ^8^ were only observed in WT.

### Peptidoglycan analysis of BWH047 and Δ*dpcA*

Based on the role of DpcA as an UndP flippase and UndP’s essential role in peptidoglycan (PG) precursor transport [52, 55], we postulated that BWH047*ΔdpcA* may have an altered PG composition compared to WT. However, liquid chromatography-mass spectrometry (LC-MS) PG composition analysis of BWH047 WT, BWH047*ΔdpcA,* and BWH047*ΔdpcA* comp strains showed identical PG profiles (**S7 Fig**). We were initially surprised by these results, but our observations are supported by recent data demonstrating that PG profiles are similar between PAO1 and PAO1Δ*dpcA* [56]. Klycheva et al. observed that loss of *dpcA* (which they refer to as *dedA4*) resulted in accumulation of the cytoplasmic PG precursor UDP-MurNAC pentapeptide but did not impact overall PG structure.

## Discussion

By combining TnSeq with *in vitro* phenotypic testing, we identified multiple intersecting pathways that are essential for extreme colistin resistance in the multidrug-resistant *Pa* clinical isolate BWH047 (**Fig 4**). We validated both previously known colistin resistance genes (e.g., *arn* operon, *pmrAB*) and identified several novel or understudied genes that contribute to colistin resistance (e.g., *dpcA*, *algU*, *wapH*). We chose to investigate one of these targets, *dpcA,* by characterizing its LPS. The lipid A of BWH047Δ*dpcA* possessed aminoarabinosylation and altered glycosyl composition and acylation status compared to BWH047 WT.

**Fig 4.**
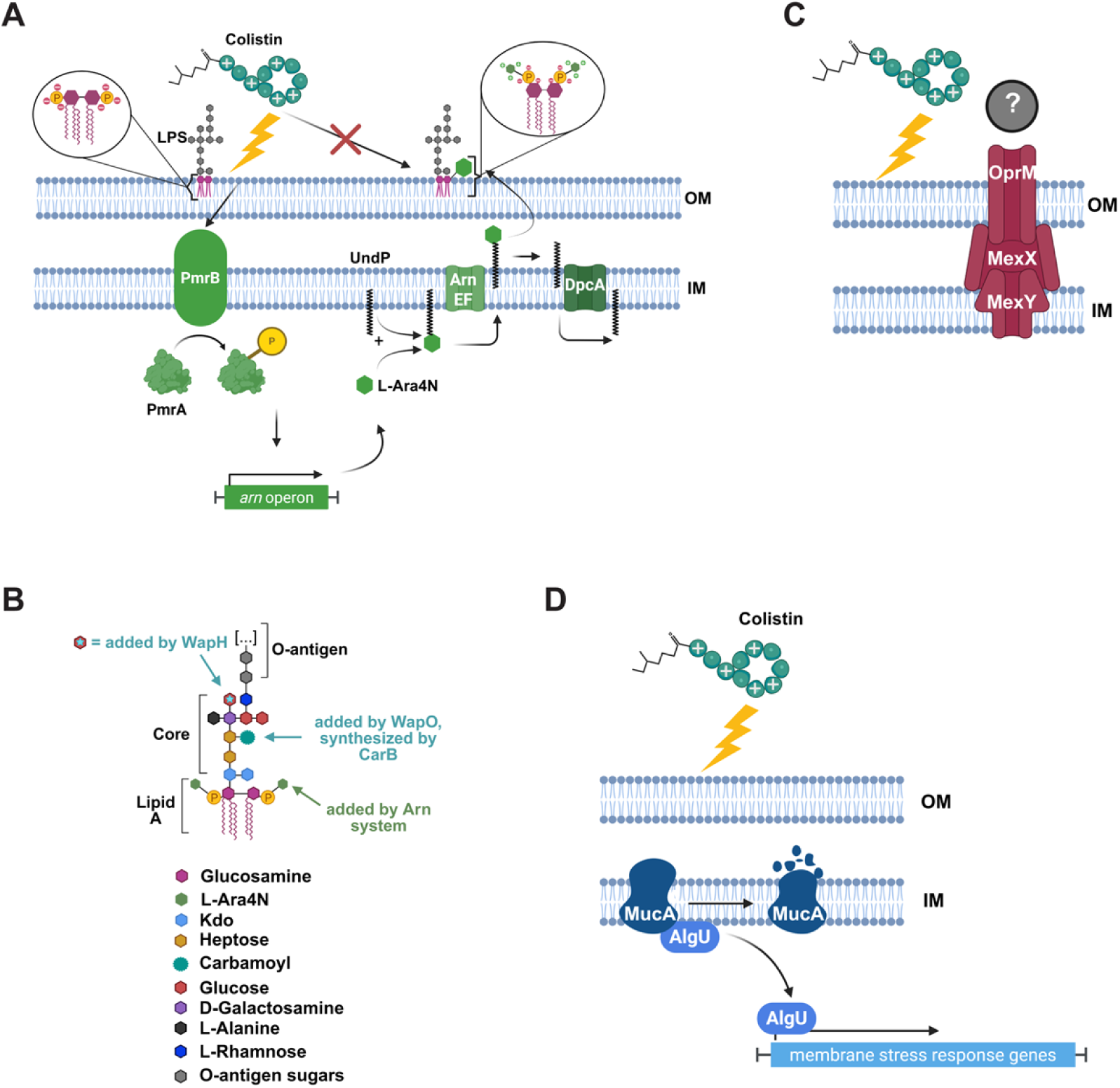
Functional model of all *Pa* BWH047 genes identified as conditionally essential for colistin resistance. (**A**) Colistin causes membrane instability due to displacement of stabilizing divalent cations. This disruption is sensed by PmrB. PmrB phosphorylates PmrA, which triggers the transcription of downstream genes including those in the *arn* operon. Following transcription of the *arn* operon, aminoarabinose (L-Ara4N) is synthesized and exported from the cytoplasm on the carrier molecule undecaprenyl phosphate (UndP) with the assistance of *arnEF* so it can be added on to lipid A. UndP is then recycled back to the cytoplasm by DpcA so it can facilitate further rounds of aminoarabinose modification. Created in BioRender. Bachta, K. (2026) https://BioRender.com/8zg018k (**B**) Detailed diagram of *Pa* LPS with the following modifications: aminoarabinose addition catalyzed by the Arn system, carbamoylization catalyzed by WapO following synthesis by CarB, and the addition of glucose (Glc) by WapH. GlcN, glucosamine; Kdo, 3-deoxy-D-*manno*-oct-2-ulosonic acid; GalN, galactosamine; L-Ala, 2-L-alanylation; L-Rha, rhamnose. Created in BioRender. Bachta, K. (2026) https://BioRender.com/cmm24hs (**C**) MexXY-OprM is conditionally essential for colistin resistance in *Pa*; however, the mechanism by which is facilitates resistance is unknown. Created in BioRender. Bachta, K. (2026) https://BioRender.com/bcq2rm0 (**D**) MucA senses membrane disruption and is then degraded to release AlgU to facilitate the transcription of membrane stress response genes; however, the mechanism by which AlgU promotes colistin resistance remains unknown. Created in BioRender. Bachta, K. (2026) https://BioRender.com/yor3ctp

In *Pa*, colistin resistance is overwhelmingly attributed to modification of the negative phosphate head groups on lipid A of LPS with the glycosyl residue, L-Ara4N [9, 12]. The products of the *arn* operon (*arnBCADTEF-ugd*) synthesize, transport, and attach L-Ara4N to lipid A [33]. In BWH047, we identified three members of this operon as conditionally essential candidates which, when deleted, validated by BMD. This suggests that lipid A modification with L-Ara4N remains the mainstay of colistin resistance even in extremely resistant *Pa* isolates (**Fig 4A**). It has been shown that transcription of the *arn* operon can be activated by five separate TCS that sense and respond to colistin or colistin-mediated changes in cation concentration [13–20]. Despite these reports, we identified conditionally essential candidate genes in only one of these systems, *pmrAB*. Intriguingly, we also identified the three candidates (*speD2*, *speE2* and *speH*) upstream of *pmrAB* (**S2C Fig**); however, deletion of any of these genes had little impact on the colistin MIC of BWH047. This is consistent with the literature which shows that deletion of *speD2, speE2,* or *speH* does not affect colistin MIC despite resulting in increased bacterial death at early time points post colistin exposure [57, 58]. Whether *speD2-E2-speH-pmrA-pmrB* are co-transcribed or expressed as two or three separate operonic units is unclear [36, 37, 59]. In BWH047, operonic prediction tools FgenesB (Softberry) and Operon-Mapper [60] predicted that all genes are located within one transcriptional unit. Therefore, the location of the *spe* genes directly upstream of *pmrAB*, their identification by TnSeq, and their lack of phenotype in our BMD assays suggests that polar effects of transposon insertion and not the direct protein products of *speD2-E2-H* are responsible for the identification of *speD2-E2-H* as gene candidates.

We also identified several candidates involved in LPS biosynthesis. This is unsurprising given that LPS, more specifically lipid A, is the target of colistin [61]. First, we identified *dnpA*, a hypothetical gene located at the end of a LPS biosynthesis operon (*wapO*-*wapH*-*mig14-dnpA*), as conditionally essential (**S2C Fig**). Surprisingly, Δ*dnpA* had a MIC unchanged from that of BWH047. Like the *speD2-E2-H-pmrA-pmrB* locus, we hypothesized that transposon insertion into *dnpA* might impact expression of other members of the operon more directly involved in colistin resistance. We were unable to delete *mig14* but successfully generated clean deletions of *wapH* and *wapO*, resulting in MICs of 1µg/mL and 160µg/mL, respectively. WapH is a hypothetical protein annotated as a putative glycosyltransferase [59]. In a 2011 study, Jochumsen et al. identified *wapH* as a putative colistin response gene but concluded that transposon insertion in *wapH* likely affected expression of the downstream *mig14* gene in a polar fashion [40]. In contrast, our results implicate *wapH* itself in colistin resistance, although we were unable to assess the role of *mig14* in colistin resistance in BWH047 because we were unable to delete it. In 2004, Matewish demonstrated that deletion of *wapH* in PAO1 resulted in truncated LPS missing portions of the outer core-OS [62]. LPS isolates from BWH047Δ*wapH* demonstrated a similar phenomenon (**S4B Fig**), which may explain the increased colistin sensitivity of this strain (**Fig 4**). The final member of this operon (**S2C Fig**), *wapO*, produces a hypothetical protein annotated as a probable carbamoyl transferase [57]. Unlike the *arn* operon, *pmrAB* and *wapH*, deletion of *wapO* in BWH047 resulted in an intermediate colistin MIC of 160µg/mL. In PAO1, mutations in *wapO* were identified as having minor contributions to extreme colistin resistance, but only in the context of prior mutations that increased the colistin MIC significantly from baseline [63]. More recently, Erdmann et al. defined adaptive mutations in *wapO* that emerged while serially passaging *Pa* in increasing concentrations of colistin [38, 39]. In PAO1, transposon insertions in *wapO* resulted in a one-fold decrease in colistin MIC (0.5 µg/mL to 0.25 μg/mL) [38, 39]. We hypothesize that the function of *wapO* may become increasingly important as colistin MIC increases, and that the low colistin MIC of PAO1 masks the ability to detect these more subtle phenotypes.

Erdmann et al. also identified adaptive mutations in *carB*, a putative carbamoyl phosphate synthase and partner of *wapO*, using experimental evolution to increase colistin resistance [38, 39]. We, too, identified *carB* as conditionally essential for colistin resistance. Whereas Erdmann et al. found that deletion of *carB* did not impact PAO1’s colistin MIC, we determined that, like Δ*wapO*, the colistin MIC of Δ*carB* was 160 μg/mL. Erdmann et al. postulated that WapO and CarB were part of a pathway responsible for carbamoylation of sugars in the LPS core-OS (**Fig 4**) [38]. Because mutations in both genes emerged during the development of colistin resistance, but complete gene deletion did not result in a change in MIC, they postulated that a decrease in carbamoylation correlated with increased colistin resistance [38, 39]. However, by utilizing BWH047, we observed the opposite—the activity of WapO and CarB, and thus the carbamoylation pathway, were required for full colistin resistance. TnSeq also identified candidates in two resistance-nodulation division (RND) efflux systems: *mexY, mexB*, and *oprM*. These genes code for members of the MexAB-OprM and MexXY-OprM complexes, which are well-characterized *Pa* tripartite efflux systems that extrude a wide variety of antimicrobials and contribute to antimicrobial resistance [41]. MexB and MexY span the inner membrane, MexA and MexX are periplasmic membrane fusion components, and OprM is the outer membrane channel protein that is shared by both efflux systems [42]. Interestingly, we found that deletion of *mexY* or *oprM*, but not *mexB,* significantly reduced the colistin MIC of BWH047. Further investigation into this phenomenon showed that *mexA*, *mexB*, and *oprM* are co-transcribed as a single transcriptional unit [64]. We hypothesize that like the *spe* system, *mexB* was identified as a candidate essential gene because of polar effects on expression of *oprM*. Thus, MexXY-OprM, but not MexAB-OprM, contributes to colistin resistance in BWH047. The literature is divided on the importance of MexAB-OprM and MexXY-OprM in *Pa* colistin resistance. Initial work to determine the substrates of these efflux pumps showed that polymyxin B, a cationic antimicrobial peptide closely related to colistin, was not a substrate for either efflux system [41]. Puja et al. showed that the MexXY-OprM system is intricately connected with *arn*- and *pmrAB*-mediated colistin resistance in *Pa,* but that this phenomenon had no impact on L-Ara4N modification of lipid A [57, 58]. It is possible that MexXY-OprM is effluxing some as-yet unidentified compound that modifies lipid A or LPS and prevents colistin’s attraction to the bacterial cell. Ultimately, how this particular efflux pump regulates colistin resistance or cross talks with the more dominant *arn-* and *pmrAB*-mediated mechanisms of colistin resistance remains unclear.

We also identified AlgU as important for colistin resistance in BWH047. Otherwise known as RpoE, AlgU is an extra-cytoplasmic sigma factor that regulates alginate production and mucoidy in *Pseudomonas* [43, 44, 65–67]. AlgU is sequestered at the inner membrane by transmembrane anti-sigma factor MucA, which is degraded through a proteolytic cascade upon sensing membrane stress [68]. AlgU also downregulates motility and virulence gene expression and upregulates stress response pathways [43, 44, 65–67]. In *Salmonella enterica* and *Escherichia coli, algU* ortholog *rpoE* was recently demonstrated to be necessary for colistin resistance [69]. Interestingly, Zeng et al. found that in *E. coli* and *S. enterica,* deletion of *rpoE* did not impact lipid A modification with L-Ara4N [69]. Instead, they attributed the colistin sensitivity of the Δ*rpoE* strain to an alteration in phospholipid composition, which resulted in decreased membrane asymmetry and increased cell permeability [69]. We have yet to determine the mechanism by which *algU* is essential for colistin resistance in *Pa*, but this remains an active area of inquiry.

Our most interesting candidate gene from TnSeq, now named *dpcA*, is orthologous to PAO1 PA4029. Deletion of *dpcA* lead to a complete loss of colistin resistance. Until recently, the function of DpcA orthologs remained unknown. Within the past few years, research in *Klebsiella pneumoniae, Burkholderia thailandensis*, *Vibrio cholerae*, and *Staphylococcus aureus* revealed that DedA is a flippase that recycles undecaprenyl phosphate (UndP) transporting it from the outer surface of the inner bacterial membrane and back into the cytoplasm [31, 45, 70–72]. In addition to transporting LPS O-antigen subunits, peptidoglycan precursors, and glycosyl residues, UndP also ferries the lipid A-modifying glycosyl residue L-Ara4N across the cytoplasmic membrane in GN bacteria [33]. Based on these studies, a model emerged wherein DedA flippases play an important role in colistin resistance by facilitating the recycling of UndP carrier molecules back into the cytoplasm once they deliver their L-Ara4N cargo to the periplasmic space. Uncharged UndP is then reloaded with L-Ara4N, facilitating further rounds of lipid A modification that lead to increasing colistin resistance (**Fig 4**) [71, 73]. In support of this model, deletion of *dedA* in either *K. pneumoniae* or *B. thailandensis* significantly decreased the amount of L-Ara4N modification of lipid A [31, 45].

Until very recently, DpcA was unstudied in *Pa*. Two papers revealed its importance for UndP recycling and implicated it as having a role in colistin resistance. A lack of DpcA led to decreased UndP recycling and diminished UndP pools on the cytoplasmic face of the bacterial inner member. Thus, fewer UndP molecules are available to transport the critical building blocks of LPS [55, 56]. However, this work was completed in the colistin-sensitive strain PAO1. We further characterized *dpcA’*s function in BWH047 by investigating LPS structure. Overall, deletion of *dpcA* had pleotropic effects on LPS including a higher proportion of core-OS sugars (Rha, Glc, Kdo, and GlcNAc) than BWH047 (**Table 2**), an increase in the content of palmitic fatty acid (16:0) and loss of the unusual cis-8-octadecanoic fatty acid (18:1Δ^8^) (**Table 3),** an increase in tetra- and hexa-acylated lipid A (**Table 6**), and a decrease in the number of L-Ara4N moieties on lipid A (**Table 5**). With respect to the LPS core-OS, UndP ferries O-antigen subunits from the cytoplasm to the periplasm [52, 53, 74]. Deletion of DpcA likely results in decreased transport of O-antigen glycosyl residues to the outer membrane leading to shorter LPS O-antigen, as observed in **S4A Fig**. GC-MS and MALDI-TOF MS/MS data revealed that the lipid A of BWH047, unlike Δ*dpcA*, contained higher levels of palmitic acid (16:0) and an unusual cis-8-octadecanoic (18:1Δ^8^) fatty acid which was totally absent in Δ*dpcA* (**Table 3**, **Fig 3, S4-S6 Figs**). While 18:1 fatty acids have been reported as a component of lipid A in *Pseudomonas syringae* and in *Francisella tularensis* [75, 76], 18:1Δ^8^ has not been reported as component of lipid A in *Pa*. The functional significance of the 18:1Δ^8^ acyl chain in lipid A has not been investigated. Future investigation will be necessary to determine biosynthetic pathways responsible for its inclusion in BWH047 lipid A, if the presence of 18:1Δ^8^ leads to increased cationic antimicrobial peptide resistance and membrane plasticity, and if and how this modification impacts lipid A recognition by the host immune system. In accordance with our hypothesis, the lipid A of Δ*dpcA* contained a lower proportion of structures that were modified by either one or two L-Ara4N moieties than in BWH047 (**Tables 2, 4, 5 and S6 Fig**). In summary, deletion of *dpcA* in BWH047 had pleiotropic and wide-ranging effects on the structure of LPS and lipid A which extend beyond its impact on L-Ara4N modification. DpcA’s role in membrane biogenesis and its relationship to colistin resistance represent an exciting area of ongoing investigation.

Overall, TnSeq identified multiple pathways conditionally essential for colistin resistance in the extremely colistin resistant clinical *Pa* isolate BWH047 and uncovered a complex web of interconnected networks that are necessary for this extreme phenotype (**Fig 4**). This work represents one of the most thoroughly validated TnSeq experiments to date. We identified 20 conditionally essential gene candidates and successfully deleted 17 target genes and 6 interesting neighboring genes. This resulted in a validation rate of 75%, wherein 15 of our 20 target genes were essential for or located in operons with genes essential for extreme colistin resistance. Taken together, TnSeq proved highly successful in identifying multiple known and novel intersecting pathways that lead to extreme colistin resistance in *Pa* and highlighted novel mechanisms that deepen our understanding of colistin resistance.

## Materials and Methods

### Ethics statement

The collection and use of *Pa* clinical isolates and associated data represented in this study was reviewed and approved by the Partners Human Research Committee under Protocol #2015P001829/BWH on September 15, 2015. The IRB waived the requirement for informed consent, as the research involved only existing bacterial isolates and retrospective clinical data.

### Bacterial strains and growth conditions

*Pa* clinical isolates were collected from Brigham and Women’s Hospital (BWH) clinical microbiology laboratory in Boston, MA between 2015 and 2016. Isolates were collected from a variety of patient sites (**Fig 1, S4 Table**) and were selected and screened for antibiotic resistance to commonly used antibiotics including aztreonam, anti-pseudomonal cephalosporins (cefepime and ceftazidime), piperacillin-tazobactam, ciprofloxacin, colistin, and meropenem. BWH047 was originally described in 2019 [77]. The remainder of the PABWH isolates are described in the present study (**Fig 1, S4 Table**). *Escherichia coli* strain TOP-10 (Invitrogen) was used for cloning, and *E. coli* strains S17.1 λpir and SM10 λpir (gift of John Mekalanos, Harvard Medical School) were used to introduce plasmids into *Pa*. Unless otherwise specified, all bacteria were grown at 37°C in LB or Mueller Hinton Broth (MHB). When necessary, antibiotics were used at the following concentrations: for *Pa*: irgasan 5 μg/mL, hygromycin 500 μg/mL and for *E. coli*, hygromycin 100 μg/mL. Additional details on strains and plasmids used in this study can be found in **S2-S4 Tables.**

### Whole genome sequencing of PABWH isolates and isogenic mutants

Briefly, genomic DNA from each PABWH isolate was harvested from a single colony and sequenced on the HiSeq 4000 Platform (Illumina, Inc., San Diego, CA) generating 2 x 150 bp paired end reads. Adapter sequences were removed, and reads were quality trimmed using Trimmomatic v.0.36 [78]. *De novo* assembly was performed using SPAdes v3.9.1 [79]. Quality control was performed by aligning trimmed reads to assembly contigs using bwa v0.7.15 [80]. All contigs shorter than 200bp or with an average fold coverage of < 5x per base were removed. To complete the draft genome sequence of BWH047 [77], Oxford Nanopore MinION sequencing was performed on a R9.4.1 flow cell after genomic DNA library preparation using the Oxford Nanopore SQK-LSK109 kit. A consensus *de novo* assembly was generated using Trycycler v.0.5.0 [81] from long read assemblies generated using Raven v.1.5.3 [82], Flye v.2.9.5 [83], and Miniasm v.0.3 [84]. The assembly was polished first with the Oxford Nanopore long reads using medaka v1.11.3 and subsequently with the Illumina short reads using polypolish v0.5.0[85] and the polca module of MaSuRCA v4.0.9 [86]. Annotation was performed using the NCBI Prokaryotic Genome Annotation Pipeline 6.10 [87].

### Whole genome phylogenetic analysis

Illumina reads from each isolate were aligned to the complete genome sequence of strain BWH047 using bwa v0.7.15 [80]. Single nucleotide variants (SNVs) were identified using bcftools v1.9 [88] using the bcftools_filter software (https://github.com/egonozer/bcftools_filter) to remove variants with SNV quality scores less than 200, read consensus less than 75%, read depths less than 5, read numbers in each direction less than 1, or locations within repetitive regions as defined by blast alignment of the reference genome sequence against itself. A maximum likelihood phylogenetic tree was generated from the core genome alignment with IQ-TREE v2.2.0 using the ModelFinder function to estimate the best-fit nucleotide substitution model by means of Bayesian information criterion (BIC) [89, 90]. Tree visualization and annotation was performed in R v4.4.2 using ggtree v3.14.0 [91].

### Generation of gene deletions and complemented strains in *Pa* BWH047

Upstream and downstream fragments surrounding the gene of interest (GOI) were amplified from BWH047 genomic DNA using primers GOI-5-1-HindIII with GOI-5-2, and GOI-3-1 with GOI-3-2-HindIII (**S1 Table**). The resulting upstream and downstream fragments, which each contained a 24-bp overlapping linker sequence (*TTCAGCATGCTTGCGGCTCGAGTT* in primers GOI-5-2 and GOI-3-1), were used as the templates for overlap extension PCR and were subsequently PCR amplified with primers GOI-5-1-HindIII and GOI-3-2-HindIII to create a single linear fragment. This was combined with the integration proficient vector pEX18Hyg [92] previously digested with HindIII (New England Biolabs) via Gibson assembly (New England Biolabs Gibson Assembly^®^ Cloning Kit) to generate allelic exchange vectors for all genes of interest (pEX18HygΔGOI, **S2 Table**). These vectors were transformed into *E. coli* TOP-10 (Invitrogen) and plated on LB agar plates supplemented with 100 µg/mL hygromycin. Plasmids were miniprepped (GeneJET Plasmid Miniprep Kit, Thermo Scientific) and verified by Sanger Sequencing (ACGT, Inc.). Vectors were transformed into *E. coli* SM10 λpir and *E. coli* S17.1 λpir. Following conjugation and allelic exchange with BWH047, gene deletions were verified using Sanger sequencing (ACGT, Inc.)

Gene deletions were complemented in one of two ways. First, *arnC* and *ugd* were complemented by re-introducing the native allele into the deletion strain at its original position in the genome. Briefly, GOI-5-1-H and GOI-2-H were used to amplify the entire WT allele. As above, Gibson assembly (New England Biolabs Gibson Assembly^®^ Cloning Kit) was used to ligate this complementation fragment into HindIII-digested pEX18Hyg generating pEX18Hyg-comp-GOI. Then, these two vectors were transformed into *E. coli* TOP-10 (Invitrogen), miniprepped (GeneJET Plasmid Miniprep Kit, Thermo Scientific), verified by Sanger Sequencing (ACGT, Inc.) and transformed into *E. coli* SM10 λpir and S17.1 λpir. Following conjugation and allelic exchange with BWH047ΔGOI, complementation was confirmed both by sanger sequencing (ACGT, Inc) and phenotypically through determination of colistin MIC (**Table 1**).

Alternatively, *dpcA* was complemented by introducing the *dpcA* sequence at the Tn7 site of BWH047Δ*dpcA* using pCOMP-hyg-npt2, a version of the pUC18T mini-Tn7T-Gm vector [93] modified with a constitutive npt2 [94] promoter and hygromycin resistance. To construct pCOMP-hyg-npt2, the vector pUC18T-mini-Tn7T-Gm-eYFP [93] was first digested with EcoRI and NsiI (NEB) and Gibson assembly was used to ligate this vector with a gene block containing a FRT-flanked hygromycin resistance cassette to generate pUC18T-mini-Tn7T_hygR_eYFP. Subsequently, pUC18TminiTN7T_hygR_eYFP was digested with BsrG1 and Kpn1 to remove the eYFP cassette. Then, Gibson assembly was used to ligate a gene block (IDT) containing both a constitutive synthetic lac promoter (PA1/04/03) [95] and a multiple cloning site was into the cut pUC18TminiTN7T_hygR to generate pCOMP-hyg. To provide alternative promoter expression, pCOMP-hyg was again digested with BstBI and KpnI and Gibson assembled with a gene block from IDT containing the npt2 promoter to generate pCOMP-hyg-npt2 (gift of J. Allen). This final plasmid was verified using whole plasmid sequencing through Plasmidsaurus (San Francisco, CA). Finally, to generate pCOMP-hyg-npt2-dpcA, the sequence of the *dpcA* was amplified from BWH047 using primers pCOMP-hyg-npt2-dpcA_5 and pCOMP-hyg-npt2-dpcA_3 (**S1 Table**) and Gibson assembly was used to ligate *dpcA* with BstBI-digested pCOMP-hyg-npt2. This final *dpcA* complementation vector was transformed into *E. coli* TOP-10 cells (Invitrogen), sequence verified via Sanger sequencing (ACGT, Inc.) and transformed into *E. coli* SM10 λpir. *E. coli* SM10 λpir containing pCOMP-hyg-npt2-dpcA, *E. coli* SM10 λpir containing pTNS2, *E. coli* HB101 containing pRK2013, and BWH047Δ*dpcA* were mated for 2 hours at 37°C [93]. Resulting colonies resistant to both hygromycin and irgasan were screened using PCR for insertion of the *dpcA* allele at the Tn7 site. Complementation was confirmed by both Sanger sequencing (ACGT, Inc.) and evaluation of colistin MIC (**Table 1**).

### Determination of minimum inhibitory concentrations (MIC)

The MICs for the PABWH collection and all BWH047 isolate derivatives were determined in triplicate in compliance with the broth microdilution (BMD) protocol as outlined by Clinical and Laboratory Standards Institute (CLSI) M100-Ed35 [25]. The following antibiotics were prepared from commercially available sources and were used to assess MICs: aztreonam, cefepime, ceftazidime, ciprofloxacin, meropenem, piperacillin-tazobactam. For most antibiotics, MICs were performed in Mueller Hinton Broth (MHB) except for colistin MICs, which were performed in cation-adjusted MHB (MHBII). *Pa* ATCC control strain 27853 was used for quality control. Colistin MICs for all PABWH isolates were initially determined using the standard colistin concentrations (0.25-128 μg/mL). Once it was established that BWH047 had a colistin MIC ≥ 128 µg/mL, colistin BMDs were performed using 10x the traditional concentrations (2.5-1280 μg/mL) to determine an exact MIC value.

### Construction of transposon mutant library in *P. aeruginosa* isolate BWH047

The pSAMHyg suicide plasmid for Himar1 mariner transposon (tn) mutagenesis [96] was transformed into *E. coli* SM10 λpir. Equal volumes of *E. coli* SM10 λpir containing pSAMHyg were mixed 1:1 with BWH047 and 125 x 10µL mate spots were spotted onto LB for 1 hour and 15 minutes. Mating conditions were optimized to ensure a minimum of 400,000 colonies of BWH047:tn were obtained. All mate spots were scraped and resuspended into a total volume of 8 mL of LB. 50 μL of this suspension was then plated onto large agar plates (n ∼ 150) supplemented with irgasan (5 μg/mL) and hygromycin (500 μg/mL) and incubated at 37°C overnight to select for colonies containing a transposon insertion event. The following day, all irgasan (5 μg/mL)/hygromycin (500 μg/mL) resistant colonies were scraped and resuspended in a total volume of 15mL of LB supplemented with 25% glycerol. The final BWH047:tn library was aliquoted and stored at −80°C for further analysis.

### Transposon Insertion (TnSeq) experimental design

An aliquot of the BWH047 transposon library was thawed and revived in MHBII at 37°C for approximately 2 hours. Following this incubation, the revived library was added to MHBII containing colistin at these final concentrations: 640 μg/mL, 320 μg/mL, and 0 μg/mL. Five experimental replicates were used per antibiotic concentration (**S2A Fig**). Cultures were incubated at 37°C with shaking for 18 hours. The colistin concentrations and library outgrowth time were optimized (1/4 MIC, 320 µg/mL and 1/2 MIC, 640 µg/mL) to provide strong selective pressure to facilitate drop out of strains with disrupted conditionally essential genes and to mirror the time point at which MICs are typically assessed (16-20 hours post inoculation) [25]. Following incubation, bacteria were harvested via centrifugation at 13,200 x *g* for 2 minutes and genomic DNA was extracted. DNA was prepared for sequencing according to the methods outlined in Kazi et al 2020 [97]. Briefly, genomic DNA was sheared to approximately 250bp (Covaris Focused Ultrasonicator). Transposon-DNA junctions were amplified, and barcodes were added using previously designed primers [97] (**S1 Table**). Library pools were quantified using Kapa library quantification kit (Roche) and Quant-iT dsDNA Assay Kit, high sensitivity (HS) (Invitrogen). Pools were sequenced on an Illumina MiSeq using the Miseq v2 reagent kit (50-cycle) generating 50bp single-end reads containing the 6bp barcode sequence, 22bp Himar transposon sequence, the TA dinucleotide insertion site, and the 22bp genome sequence flanking the TA site. Illumina reads were processed with TIS_Tools (github: https://github.com/egonozer/TIS_tools). Briefly, the transposon sequence was identified in each read, allowing for up to a 1 base mismatch. Next, reads were trimmed to a final length of 24bp containing only the TA dinucleotide insertion site and the 22bp of BWH047 genome sequence that flanked the TA dinucleotide. Then, the number of sequences for each TA insertion site was counted and grouped within each library and reads were mapped to the BWH047 complete genome sequence using Bowtie v1.3.1 [98]. Reads were excluded from analysis if they aligned to ≥ 1 genomic site or contained > 1 base mismatch. Finally, the alignment position and strand (+ or -) were assigned before outputting a file associated with each experimental replicate. The filtering done to determine high confidence TA insertion sites, the percentage of reads retained for analysis after each filter was applied (separated by exposure condition—320 μg/mL, 640 μg/mL, and MHB II only), and the percentage of TA sites included in the final analysis are depicted in **S3 Fig**.

### TnSeq data analysis and identification of conditionally essential genes

To identify genes conditionally essential for colistin resistance in BWH047, a modified version of the ESSENTIALS pipeline [99] was implemented (github: https://github.com/egonozer/essentials_local). Briefly, with read aligning to flanking sequences in the reference BWH047 sequence were identified, and sites which had a combined total of < 3 aligned in any of the control libraries (MHBII only, 0 µg/mL) were filtered out. Additionally, any TA sites located in the last ∼ 5% of a protein-coding sequence were excluded from analysis. Next, a Loess normalization was executed to reduce the bias of higher read counts near the chromosomal origin of replication given its enhanced TA site accessibility during chromosomal replication. Finally, a Fisher’s exact test followed by Benjamini-Hochberg correction was employed to determine which genes were conditionally essential for survival in the presence of colistin. We defined a gene as conditionally essential if it met the following criteria: log_2_ fold change < −2.0 and false discovery rate (FDR) < 0.05 when comparing transposon insertion count per gene when grown in the presence of either 640 μg/mL or 320 μg/mL colistin treatment (treatment) compared to transposon insertion count when grown in MHBII media alone (control, 0 µg/mL).

### Extraction and purification of *Pa* LPS

Twenty liters of BWH047 and BWH047Δ*dpcA* were grown in LB to exponential phase (OD∼0.8), at which point cultures were fixed in 1% phenol for 1 hour and then pelleted at 8,000 x *g* for 20 minutes at 4°C. LPS was extracted from cell pellets by adding an equal volume of preheated 90% liquefied phenol and gently stirring for 20 minutes at 68°C, according to the Westphal extraction procedure [51]. After LPS extraction, the mixture was cooled in an ice bath and centrifuged at 10,000 x *g* for 20 minutes at 4°C. The water layer was collected, and the remaining phenol phase was extracted an additional two times with water, following the same protocol as above. The combined extracts of all three water layers and the residual phenol layer were extensively dialyzed against deionized water (12-14,000 MWCO), freeze-dried, and washed with 9:1 (v/v) ethanol in water at 4°C. Nucleic acids and proteins were removed by treating overnight with Benzonase (37°C), followed by proteinase K (37 °C) digestion (50 mM MgCl_2_•6H_2_O and 20 mM NaOAc•3H_2_O) and dialysis (12-14,000 MWCO) at 4°C against several exchanges of dH_2_O. Dialyzed phenol and water phases were then freeze-dried, and the LPS preparations were suspended in deionized water. These LPS fractions were further purified by ultracentrifugation at 100,000 × *g* at 4°C for 16 h. Enzymatic purification steps and ultracentrifugation were repeated twice to prepare ultrapure LPS. In addition, purified LPS was deprived of phospholipids and free fatty acids using 90% EtOH washes (completed three times) followed by short sonication at 0°C and collection of LPS by centrifugation at 3,500 x *g*. A trace of coextracted phospholipids was removed from the final LPS preparation using three 90% EtOH washes, facilitated by short sonication at 0°C and centrifugation at 3,500 x *g*. The ultrapure LPS was used for all chemical studies.

### DOC-PAGE analysis of LPS

Purified *Salmonella typhimurium*, *P. aeruginosa* BWH047, BWH047Δ*dpcA*, and BWH047Δ*wapH* LPS were resolved on a 18% polyacrylamide gel (PAGE) containing deoxycholic acid (DOC) detergent[100]. The gel was developed using a silver stain reagent kit (Bio-Rad) after oxidation with sodium periodate.

### Chemical composition analysis of LPS

A simultaneous analysis of glycosyl residues and fatty acids constituting BWH047 and BWH047Δ*dpcA* LPS was achieved through sample derivatization to O-trimethylsilyl (TMS) methyl ethers for sugar residues and fatty acid methyl esters (FAME) and TMS-FAME derivatives for hydroxylated fatty acids. Briefly, samples were methanolyzed with 1M HCl-methanol at 80°C for 18 h, re-N-acetylated at 100°C for 1 h, and O-trimethylsilylated with Tri-Sil reagent at 80°C for 30 min. Each sample was supplemented with myoinositol, which was used as an internal standard [101, 102].

The analysis of TMS and FAME derivatives was performed by gas chromatography-mass spectrometry (GC-MS) on an Equity-1 (Supelco) fused silica capillary column (30 m length × 0.25 mm ID × 0.25 μm film thickness) interfaced with a Hewlett-Packard HP5890 gas chromatograph equipped with a mass selective detector 5970 MSD. The oven temperature was initially set at 80°C for 2 min, then increased to 140°C at a rate of 20°C/min with a 2 min hold and then raised to 200°C at a rate of 2°C/min, followed by an increase to 250°C at a rate of 30°C/min with a 5 min hold. The data were processed using Agilent ChemStation.

### Isolation of lipid A from intact LPS

To prevent the loss of acid-labile L-Ara4N and phosphate residues, ultrapure LPS from *Pa* BWH047 and Δ*dpcA* was dissolved in 10 mM sodium acetate pH 4.5 for 1h at 100°C with gentle agitation. Lipid A was recovered from the sodium acetate solution by threefold extraction with chloroform and, finally, the extraction of the combined chloroform phases with water. The remaining chloroform phase was dried, and purified lipid A was used for MALDI-TOF MS analysis.

### MALDI-TOF MS and MS/MS analysis of lipid A

Approximately 2 µL of purified *Pa* BWH047 and Δ*dpcA* lipid A dissolved in chloroform:methanol (3:1, v/v) was mixed with 0.5 M 2,4,6-trihydroxyacetophenone matrix (Sigma) in methanol at a 1:1 ratio (v/v), and 0.5-1 µL was immediately applied to a Matrix-Assisted Laser Desorption/Ionization (MALDI) plate using a microcapillary. Spectra were acquired on a MALDI-TOF (time of flight) mass spectrometer RapifleX TissueTyper system (Bruker Daltonics, Billerica, MA, USA) in negative reflector ionization mode [M-H]^-^. From select observed precursor ions, MS/MS tandem mass spectrometry was performed on the same instrument in TOF/TOF geometry using a single-beam laser and nitrogen collision-induced dissociation (CID) in the negative ionization mode, with an isolation window of ±10MhZ and 4000 fragment shots. All data were processed using flexAnalysis (v. 4.2) MS software (Bruker Daltonik, GmbH).

### Comparative analysis of LPS and lipid A composition

Monosaccharide composition of LPS isolated from each strain (**Table 2**) was determined by calculating the relative molar percentage of each monosaccharide from response factor–corrected values based on the response factor of original monosaccharide standards GC–MS (**S4 Fig**). Due to the unavailability of authentic standards for FucNAc, Ara4N, and QuiNAc, we used the relative response factor of GlcNAc. Fatty acid composition of LPS isolated from each strain (**Table 3**) was calculated using the relative response from the GC-MS EI detector, where the sum of all integrated fatty acid peak areas equals 100%. The peak intensities (**S8 Table**) for all peaks with assigned proposed structures of interest (**Table 4**, **S6 Fig**) were divided by the sum of all peak intensities with proposed structures to determine a relative percentage of various glycosyl residues, acylation patterns, and L-Ara4N in each sample with the indicated characteristic.

### Peptidoglycan purification and analysis

Peptidoglycan (PG) was isolated and purified from 350 mL of mid-log phase cultures using previously published procedures [103–105]. BWH047, BWH047Δ*dpcA*, and BWH047Δ*dpcA*-comp were grown overnight, back-diluted 1:100 into 350 mL of fresh LB, and cultured to mid-log exponential growth phase. Cells were then pelleted, washed twice with PBS, and frozen at −80°C until PG purification.

To isolate PG sacculi, the washed cell pellets were thawed on ice, resuspended in cold PBS, and added dropwise to boiling 10% SDS (5% SDS final). After 1 hour, the insoluble material was collected by ultracentrifugation (242,500 x g, 60 min, 30°C) and the resulting pellets were washed five times with warm (40°C) autoclaved DiH2O. Intact peptidoglycan sacculi were treated with chymotrypsin (0.3 mg/mL) overnight with shaking (540 rpm) at 37°C to remove bound proteins. Excess chymotrypsin was denatured, and cleaved material solubilized, by adding 0.5% SDS (final, v/v) and incubating each preparation for 30 mins at 80°C. Samples were washed three times using warm (40°C) autoclaved DiH2O using ultracentrifugation, as described above. Soluble muropeptides were produced by incubating intact chymotrypsin-treated sacculi with mutanolysin (62 U/mL final) and 5μM NaHPO4/NaH2PO4, pH 5.5 overnight with shaking (540 rpm) at 37°C. Afterwards, undigested material was re-treated with mutanolysin, as described above, and incubated for an additional 4 to 6 hours. Mutanolysin was heat-inactivated (10 min, 100°C) and soluble muropeptides were harvested via centrifugation at 22,000 x g for 30 min. Isolated supernatant containing soluble muropeptide fragments was frozen (−80°C) and dried (FreeZone Benchtop Freeze Drier, LABCONCO). The purified muropeptides were resuspended in 150μL of saturated 0.5M sodium borate buffer, pH 9.25. A 50 μL aliquot of 100 mg/mL NaBH4 was added dropwise to reduce each sample and, after 1 hr incubation, the reactions were quenched with approximately 10μL of formic acid (final pH 3). The samples were then frozen and lyophilized as described above. Reduced muropeptides were resuspended in H2O:MeOH (200 μL, 9:1, v:v) containing 0.1% formic acid and sonicated in a water bath (Fisher Scientific) for 10 min. Soluble reduced muropeptides were collected by transferring 180μL of supernatant into an LC-MS vial after centrifugation at 13,000 x g at 4°C for 15 min. Samples were then stored at −20°C until LC-MS analysis.

### LC-MS analysis of muropeptides

Analyses were performed as published previously on a Shimadzu LCMS9030 QToF system coupled to a LC-40B X3 UPLC, a SIL-40C X3 autosampler (10°C), and a CTO-40C column oven (40°C) [104, 105]. Borohydride-reduced mutanolysin-digested muropeptides were separated on a Waters BEH C18 column (2.1 x 50 mm, 1.7μm particle size), at a flow rate of 0.3 mL/min. Solvent A was water and Solvent B was MeOH, with both solvents containing 0.1% formic acid (v/v). The initial solvent condition was 99:1 (A:B) for 3 min. This was followed by stepwise linear gradients first to 8% Solvent B (12 min), then to 20% Solvent B (24 min), and finally 95% Solvent B (25 min), followed by a hold at 95% Solvent B until 29 min. The column was then re-equilibrated to 1% Solvent B (1 min of 1% Solvent B, followed by a 5 min re-equilibration time). The reduced muropeptides were subjected to electrospray ionization in positive ion mode using data-dependent acquisition. Interface voltage was 4.50 kV at 300°C with a desolvation temperature of 526°C and a DL temperature of 250°C. Gas flows for nebulizing, heating, and drying gases were 2, 10, and 10 L/min, respectively. The m/z range for the precursor MS scan was 100–2000.

## Acknowledgments

We thank the Brigham and Women’s Hospital (BWH) clinical microbiology laboratory for assistance in collecting *Pa* isolates and Dr. Francisco Marty for assistance in coordinating the collection. We would like to thank the Allen laboratory at Loyola University Stritch School of Medicine for providing the pCOMP-hyg cloning vector and a protocol for its use. We acknowledge Griffin Keiser (CCRC) for his help with the initial steps of LPS purification. We would like to thank all members of the Hauser, Ozer and Allen laboratories for their valuable comments during numerous discussions of this work.

## SUPPORTING INFORMATION CAPTIONS

**S1 Table.** List of primers used in this study.

**S2 Table.** List of plasmids used in this study.

**S3 Table.** List of strains in this study.

**S4 Table.** List of PABWH Collection Isolates and associated metadata. Contains source, sequence type, T3SS genotype, MIC, and SIR data.

**S5 Table.** ESSENTIALS bioinformatic pipeline output from transposon insertion sequencing experiment exposing BWH047 to 320 µg/mL of colistin.

**S6 Table.** ESSENTIALS bioinformatic pipeline output from transposon insertion sequencing experiment exposing BWH047 to 640 µg/mL of colistin.

**S7 Table. Gene targets from the transposon insertion experiment that passed the filtering conditions for both 320 µg/mL and 640 µg/mL colistin.**

**S8 Table.** MALDI-TOF MS raw signal integration values for BWH047 and BWH047 Δ*dpcA* lipid A released using a mild procedure (pH 4.5).

**S9 Table.** TIS_tools bioinformatic output for TnSeq read processing.

**S10 Table.** TIS_tools bioinformatic output for TnSeq TA site analysis.

**S11 Table.** Muropeptide analysis of *P. aeruginosa* BWH047 and its derivatives. Includes raw peak areas and percentage of total peak areas for both replicates in all strains.

**S1 Fig. Antibiotic resistance profiles of the PABWH P. aeruginosa isolate collection**. Minimum inhibitory concentrations as determined by BMD for aztreonam (**A**), cefepime (**B**), ceftazidime (**C**), ciprofloxacin (**D**), meropenem (**E**), and piperacillin-tazobactam (**F**) for all isolates within the PABWH collection (n = 85). The x-axis is the concentration of antibiotic in µg/mL and the y-axis is total number of isolates with each MIC. MICs are reported as susceptible (S, green), intermediate (I, yellow), or resistant (R, red), based on current CLSI guidelines (Ed. 100, M35). (**H**). Summary antibiogram for all 8 antibiotics assayed by broth microdilution for the PABWH isolate collection including colistin. The y-axis represents the percent of isolates that are sensitive (green), intermediate (yellow) or resistant (red) for each antibiotic list on the x-axis.

**S2 Fig.** TnSeq Experimental Design, Transposon Insertion Density Map, and Operonic Diagrams of TnSeq Hits. (A) Experimental design of TnSeq experiment challenging BWH047 transposon library with ¼ and ½ MIC colistin. Created in BioRender. Bachta, K. (2026) https://BioRender.com/os150y1 (**B**) Linearized heatmap showing distribution of transposon insertion sites across the BWH047 6.86Mbp genome with sites with greater frequency of insertions present as yellow and areas of lower density transposon insertions present as blue. The black region around 2.5Mbp represents a large ∼90kbp region containing a duplication for which transposon depth was unable to be evaluated because the genes are diploid. Insertions per site is calculated in 1,000bp ranges. (**C**) Operonic context for TnSeq hits with validated genes or validated neighboring genes. Minimum inhibitory concentrations of individual gene deletions indicated underneath. TnSeq gene hits from original screen indicated with *; – indicates gene deletion was not pursued. Created in BioRender. Bachta, K. (2026) https://BioRender.com/ggazop0

**S3 Fig. Quality Control for the creation of BWH047 transposon library and experimental execution**. (**A**) Filtering schematic depicting methodology for determining high-confidence, analyzable TA sites present in the MHB II media control pools (n=5). Created in BioRender. Bachta, K. (2026) https://BioRender.com/o0pjlgr (**B**) Mean percentage read retention ± 1 SD from TnSeq by experimental condition. For each of the three experimental conditions (320 and 640 μg/mL colistin and MHBII), the mean percentage is calculated from 5 replicates. (**C**) Mean percentage TA site occupancy by experimental condition ± 1 SD. For each of the three experimental conditions (320 and 640 μg/mL colistin and MHBII), the mean percentage is calculated from 5 replicates, and the denominator is the total number of genomic TA sites (n=107,934). Analyzable TA sites (n=16,913) are defined as TA sites present in all 5 MHB II media control pools with ≥ 3 reads in at least one pool. All data presented in this figure represents TIS_tools preprocessing outputs and is shown in Supplementary Tables 10 and 11.

**S4 Fig. Comparative analysis of LPS from selected strains of *P. aeruginosa* BWH047.** The purified LPS was resolved on PAGE and visualized with silver after periodate oxidation. (**A**) LPS isolated from *Salmonella typhimurium* (control), *P. aeruginosa* BWH047, Δ*dpcA*, and (**B**) Δ*wapH*. Extracted LPS from both *P. aeruginosa* BWH047 WT (**C**) and Δ*dpcA* (**D**) was further hydrolyzed with 1M HCl-MeOH, re-*N*-acetylated, and the resulting monosaccharides and fatty acid residues were converted to TMS-methyl glycosides and TMS-fatty acid methyl esters (FAME-TMS, for hydroxyl fatty acids) and fatty acid methyl esters (FAME, for non-hydroxylated fatty acids) and analyzed by GC-MS. The summary of the glycosyl composition is listed in **Table 2**, and of FAME/FAME-TMS in **Table 3**. Rha, rhamnose; Ara4N, 4-amino-4-deoxy-L-arabinose; FucNAc, N-acetyl fucosamine; Man, mannose; Kdo, 3-deoxy-D-manno-oct-2-ulosonic-acid; QuiNAc, N-acetylquinovosamine; Glc, glucose; GalNAc, N-acetylglucosamine; GalNAc, N-acetylgalacatosamine; Ino, inositol; 10:0, decanoic acid; 10:0(3-OH), 3-hydroxydecanoic acid; 10:0(2-OH), 2-hydroxydecanoic acid, 10:2(3-OH), 3-hydroxydecadienoic acid; 16:0, hyexadecanoic (palmitic) acid; Δ^8^, cis-8cis-8-octadecanoic acid.

**S5 Fig. Collision-Induced Dissociation (CID) MS/MS analysis of selected ions corresponding to lipid A substituted with 18:1, cis-8-octadecanoic acid.** (**A**) Parent ion at m/z 1404 (Peak I) was fragmented in GC MS/MS. Detected fragment ions (peaks corresponding to fragments from proposed structures) are presented in bolded red alongside corresponding portions on the proposed structures (dashed lines, inset). (**B**) Parent ions at peaks F’ and F at m/z 1375.9 and 1391.9, respectively, were fragmented using GC MS/MS. Detected fragment ions (peaks corresponding to fragments from proposed structures) are presented in bolded red alongside corresponding portions on the proposed structures (dashed lines, inset). Peak F’ (m/z 1375.9) and Peak F (m/z 1391.9) are related by the addition of an –OH group, so only one probable structure is depicted (**B/C**, inset) representing both peaks with the aforementioned –OH group depicted in blue.

**S6 Fig. Structural heterogeneity of *P. aeruginosa* BWH047 lipid A revealed by MALDI-TOF-MS.** The listed, proposed lipid A structures (A-U) correspond to the major ions detected during MS analysis (Fig 3 and Table 4) and highlight the proposed structures of lipid A. Structures and m/z values marked with (‘) indicate a different structural version of lipid A deprived of hydroxyl group (-OH) on the acyloxy acyl chain. “Na” indicates the presence of a sodiated version of lipid A.

**S7 Fig. DpcA does not alter peptidoglycan composition. (A)** Peptidoglycan was purified from biological replicate cultures, digested with mutanloysin, and the muropeptide profile was analyzed by LC-MS. Strains included BWH047 (WT, black), Δ*dpcA* (blue), and complement (*dpcA* comp, purple). (**B**) Quantification of relative peak area from data collected in ‘A’. No statistical significance observed between strains using 2-Way ANOVA (*p* > 0.05) with multiple comparisons and Tukey post-hoc test.

